# Large-scale differences in functional organization of left- and right-handed individuals using whole-brain, data-driven analysis of connectivity

**DOI:** 10.1101/2021.08.28.458027

**Authors:** Link Tejavibulya, Hannah Peterson, Abigail Greene, Siyuan Gao, Max Rolison, Stephanie Noble, Dustin Scheinost

## Abstract

Handedness influences differences in lateralization of language areas as well as dominance of motor and somatosensory cortices. However, differences in whole brain functional organization due to handedness have been relatively understudied beyond pre-specified networks of interest. Functional connectivity offers the ability to unravel differences in the functional organization of the whole brain. Here, we compared connectivity profiles of left- and right-handed individuals using data-driven parcellations of the whole brain. We explored differences in connectivity profiles of previously established regions of interest, and showed functional organization differences between primarily left- and primarily right-handed individuals in the motor, somatosensory, and language areas using functional connectivity. We then proceeded to investigate these differences in the whole brain and found that the functional organization of left- and right-handed individuals are not specific to regions of interest. In particular, we found that connections between and within-hemispheres and the cerebellum show distinct patterns of connectivity. Together these results shed light on regions of the brain beyond those traditionally explored that contribute to differences in the functional organization of left- and right-handed individuals.

## Introduction

Left-handed individuals comprise approximately 10% of the population^1^. This rare event in the population is believed to be due to the development of language lateralization in the left hemisphere, giving rise to a primarily right-handed population^2^. As such, the association between language lateralization and handedness have been well studied^3, 4^. Additionally, brain differences between left- and right-handed individuals extend beyond language lateralization including differences in the motor and the somatosensory networks^5, 6^. Neuroimaging studies have begun to highlight these differences using both functional activation^7–10^ and morphometry^11–13^. Even so, these types of studies do not address how brain regions interact and, therefore, may give an incomplete picture of the brain correlates of handedness. Functional connectivity analysis using functional magnetic resonance imaging (fMRI) is a powerful tool to characterize group differences exhibiting temporal synchrony of activity among brain regions.

While there have been some functional connectivity studies of handedness^14–16^, these are limited to specific networks chosen *a priori* and potentially fail to capture a complete picture of the connectivity profiles of handedness^7–10^.

In this study, we utilize resting-state fMRI data, functional connectivity analyses, and cluster- based inference^17^ to identify differences between left- and right-handed individuals using both hypothesis-based (e.g., networks of interest) and data-driven (e.g., whole-brain) approaches across two large datasets. For the hypothesis-based analyses, we define *a priori* networks of interest based on previous literature to investigate connectivity differences in the motor^18^, somatosensory^6^, and language^19, 20^ networks. For the data-driven analyses, we calculate whole-brain functional connectomes (i.e., a functional connectivity matrix containing pair-wise connections from all brain regions) using a 268-node functional brain parcellation^21^. As handedness preferences are well-established by 5 years of age^22^, we chose to investigate connectivity differences between left- and right-handed individuals using data from two developmental datasets^23, 24^, the Healthy Brain Network (HBN) and the Philadelphia Neurodevelopmental Cohort (PNC). Because our sample contains school-aged children, adolescents, and young adults, we can better investigate innate differences in functional organization as opposed to the effects of adaptive differences caused by left-handed individuals interacting in environments typically designed for right-handed individuals.

First, we performed cluster-based inference on our primary dataset, the HBN, establishing robust patterns of connectivity differences in the motor, somatosensory, and language networks. We then estimated the generalizability of these results to the PNC. Given the consistency of results and to increase power for whole-brain analyses, we combined these datasets to examine differences across the connectome and perform exploratory investigations of differences for within- and between-hemispheric edges, and cerebellar edges. Overall, these results demonstrate that wide-spread differences in functional organization, spanning the whole-brain, exist between left-handed and right-handed individuals. Thus, it may be important to account for handedness in functional connectivity studies, in particular for studies involving neuropsychiatric disorders, where left-handed individuals are disproportionately represented^25^.

## Results

Data obtained from the Healthy Brain Network (HBN)^23^ were used for the networks of interest results. These networks of interest are based on previous literature that has shown functional differences between left- and right-handed individuals. After excluding subjects for missing data and excessive motion (>0.2mm), 905 individuals remain (right-handed: 787, left-handed: 118), with ages ranging from 5-21 years. Edinburgh Handedness Questionnaire (EHQ) scores were used as a measure of the extent individuals were left-handed and right-handed. For generalization of networks of interest results, we used data from the Philadelphia Neurodevelopmental Cohort (PNC)^24, 26^. After excluding subjects for missing data and excessive motion (>0.2mm), 859 subjects remain (right-handed: 742, left-handed: 117) with ages ranging from 8-21 years. Measures of handedness were based on self-report of dominant hand in order to complete a finger tapping task.

Resting-state fMRI data from both datasets were processed with identical, validated pipelines and parcellated into 268 nodes (the Shen atlas) using a whole-brain, functional atlas defined previously in a separate sample.^21^ Next, the mean time courses of each node pair were correlated and correlation coefficients were Fisher transformed, generating a connectome for each participant. These connectomes were subsequently used in cluster-based inference, either restricted to *a priori* networks of interest (i.e., motor, somatosensory, and language) or at the whole-brain level. To define our networks of interest, we translated Brodmann areas from previous literature onto the Shen atlas nodes (Table 1), visualizations of node allocations shown in (Fig. S11). Handedness from the HBN was analyzed using EHQ scores as both continuous values (when analyzing data from HBN independently) and as dichotomized data (when combined with the PNC for data harmonization purposes). Only binary handedness data from the PNC was available. Analyses were performed using the network-based statistic (NBS), specifying a target familywise error rate of 5%.

**Table 1:** Allocation of nodes for each of the three networks of interest: motor, somatosensory, language. Node definitions for both left and right hemispheres are based on Brodmann Areas as reported from previous literature of differences in pure activation patterns.

| Network | Brodmann Areas | Shen atlas nodes |  |
| --- | --- | --- | --- |
|  |  | Left | Right |
| Motor | Premotor/ Supp. motor: BA6<sup>27</sup><br>Primary Motor: BA4<sup>28</sup> | BA4: Node 158<br>BA6: Node 157, 159-166, 218 | BA4: Node 23<br>BA6: Node 24, 26, 27, 29-32 |
| Somatosensory | Primary Sensory: BA1<sup>29</sup> | BA1: Node 167, 171-173 | BA1: Node 33, 38-40 |
| Language: Broca's, Wernicke's | BA 44, BA9, BA 40<sup>8</sup><br>Wernicke's: BA22<sup>30</sup>, BA39<sup>31</sup> | BA9: Node 145-147<br>BA22: Node 197<br>BA39: Node 182-184 | BA9: Node 10, 11<br>BA22: Node 63, 64<br>BA39: Node |
|  | • Broca's: BA44, BA45<sup>32</sup> | • BA40: Node 179-181<br>• BA44: Node 156<br>• BA45: Node 155 | 48, 49<br>• BA40: Node 45-47<br>• BA44: Node 21, 22<br>• BA45: Node 20 |

### Networks of Interest in HBN datasets

First, we examined the motor, somatosensory, and language networks in the HBN using EHQ as a continuous measure of handedness, where scores ranged from -100 (extremely left-handed) to 100 (extremely right-handed). The network-based statistic^17^ was used to identify edges in a connectome that are significantly different between groups of individuals that are primarily left-handed (left- > right-handed) and primarily right-handed (right- > left-handed).

#### Motor

Within the motor network (Fig. 1A/S1A), two clusters consisting of 227 edges (right- > left-handed) and 195 edges (left- > right-handed) show significantly different (p<0.05, two-tailed, corrected) connectivity between groups. Interhemispheric connections between both sides of the motor strip exhibited a mix of greater and weaker connectivity for the left- > right-handed group compared to right- > left-handed group. However, edges between the motor areas and other regions of the brain show distinct patterns between the left- > right-handed group and right- > left-handed group. In the right- > left-handed group, edges of greater connectivity compared to left-handed individuals are scattered throughout the brain across all anatomical regions. Notably, the majority of these edges are between-hemisphere edges relative to within-hemispheres (between: (136/227 edges or 59.9%; within: 91/227 edges or 40.0%; χ^2^=4.31, p=0.038; Fig. S2).

**Fig. 1:**
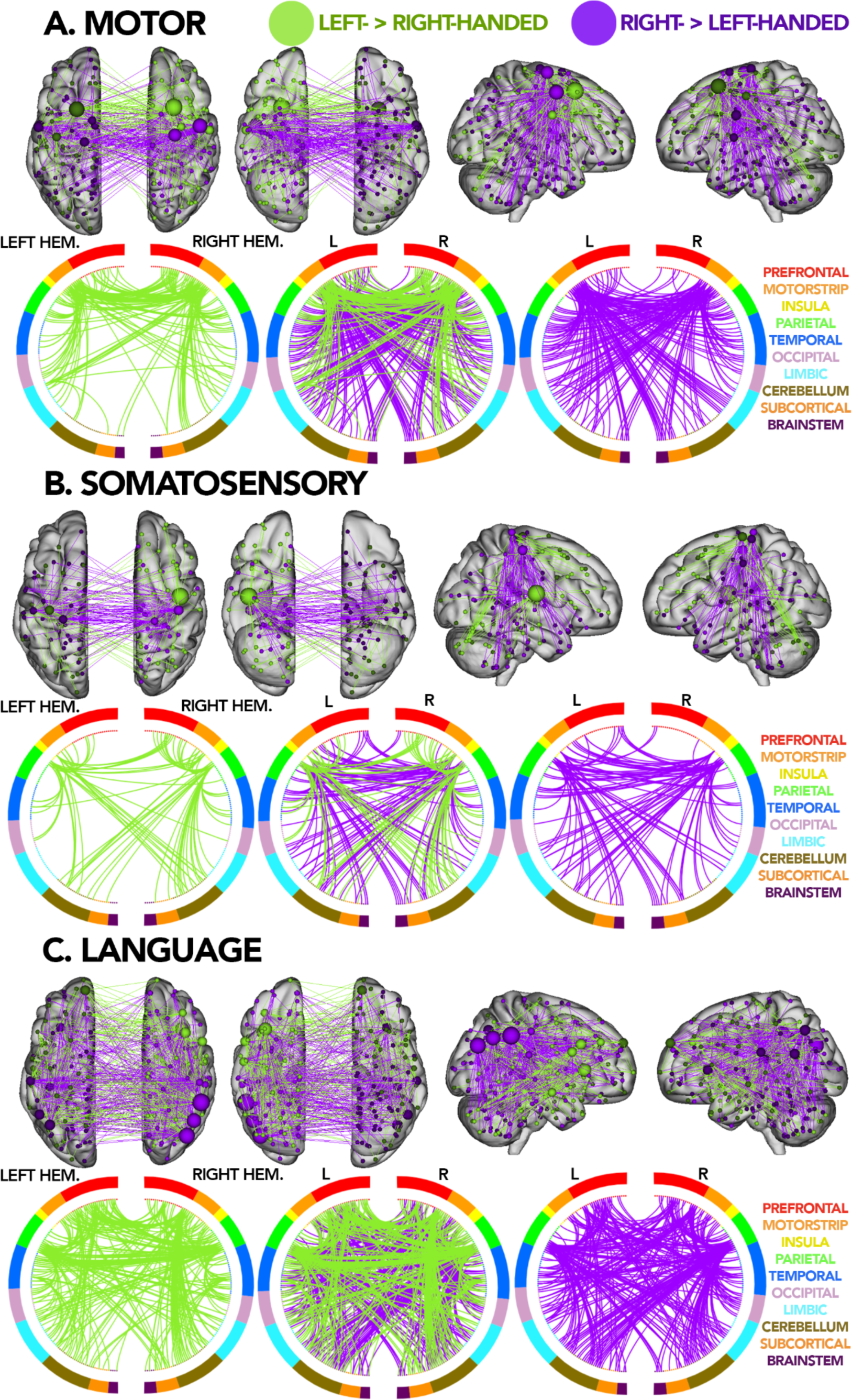
Brain and circle plots for each of the a priori defined networks in the HBN dataset. Edges that are greater for the left- > right-handed group are shown in green while edges that are greater for the right > left-handed group are shown in purple. Top row for each section shows significant edges drawn on an anatomical 3D brain with nodes sized based on the number of significant edges identified. Bottom row for each section shows circle plots where the left and right hemispheres are depicted as left and right semi-circles, respectively. The middle circle plot shows an overlay between left- and right-handed individuals. Nodes are color-coded by anatomical region constructed based on the Shen atlas, each line depicts a significant edge identified through NBS. Legend for which anatomical region each color represents is shown next to the circle plots. Each section shows results for each network of interest: (A) motor (p-val: 0.027), (B) somatosensory (p-val: 0.024), (C) language (p-val: 0.005).

In contrast, a majority of edges showing greater in the left-> right-handed groups are within-hemisphere edges with between-hemisphere generally being confined to motor-motor edges (between: 82/195 edges or 42.0%; within: 113/195 edges or 58.0%; χ^2^=2.64, p=0.104; Fig. S2). Neither group exhibit edges confined to a specific hemisphere (left- > right-handed: χ^2^=1.79, p=0.181; right- > left-handed: χ^2^=1.42, p=0.233; Fig. S2). Perhaps most interestingly, we observe a bundle of edges between the right motorstrip and the ipsilateral cerebellum in left-handed individuals, in alignment with the known roles for the cerebellum in motor control and adjustments^33^. Yet, canonical motor-cerebellar circuits point towards contralateral connections (i.e., the right motor strip connects to the left cerebellum). That we observed stronger ipsilateral functional connections in left-handed individuals may point toward neuroplasticity of left-handed individuals needing to adapt to a primarily right-dominant society (e.g., scissors, computer mice)^34^.

#### Somatosensory

Similar patterns are observed for the somatosensory network (Fig. 1B/S1B) with two clusters consisting of 127 edges (right- > left-handed) and 88 edges (left- > right-handed) exhibiting significantly different (p<0.05, corrected) connectivity between left- > right-handed and right- > left-handed groups. Neither group exhibit edges lateralized to specific hemisphere (left-> right-handed: χ^2^=2.22, p=0.136; right- > left-handed: χ^2^=0.04, p=0.841; Fig. S2). For edges of greater connectivity in right-handed individuals, a majority are between-hemisphere edges relative to within-hemispheres (between: 80/127 edges or 63.0%; within: 37/127 edges or 37.0%; χ^2^=7.81, p=0.005; Fig. S2).

For edges of greater connectivity in the left- > right-handed group, a majority are within-hemisphere edges relative to between-hemispheres (between: 29/88 edges or 33.0%; within: 59/88 edges or 67.0%; χ^2^=5.27, p=0.022; Fig. S2). However, of the contralateral edges identified as greater connectivity in the left- > right-handed group, the majority are edges stemming from the parietal networks to the contralateral cerebellum on both sides This could be partly due to the previous phenomena explained in the motor network. Studies have also shown that the cerebellum has representation of somatosensory^35^, accounting for the synchronous activity observed in both populations, but particularly in left-handed individuals.

#### Language

Finally, for the language network (Fig. 1C/S1C), two clusters consisting of 337 edges (right- > left-handed) and 325 edges (left- > right-handed) display significantly different (p<0.05, corrected) connectivity between left- > right-handed and right- > left-handed groups. Similarly to the patterns observed in the motor and somatosensory networks, the connectivity of cerebellum is notable. Bundles of edges, exhibiting greater connectivity in the left- > right-handed group, between both frontal lobes to the ipsilateral cerebellum are present. Additionally, bundles of edges with greater connectivity in right-handed individuals are observable between nodes in the right parietal lobe and both hemispheres of the cerebellum.

In contrast to the motor and somatosensory networks, no differences in the distribution of between and within-hemisphere edges in the right- > left-handed group is observed (between: 165/337 edges or 49.0%; within: 172/337 edges or 51.0%; χ^2^=0.05, p=0.823; Fig. S2).

However, similar to the motor and somatosensory networks, a majority of edges exhibiting greater connectivity in the left- > right-handed group are within-hemisphere edges relative to between-hemispheres (between: 133/325 edges or 40.9%; within: 192/325 edges or 59.1%; χ^2^=5.22, p=0.022; Fig. S2) with between-hemisphere edges primary located between the parietal lobes. Neither group exhibit edges lateralized to specific hemisphere (left-handed: χ^2^=3.40, p=0.065; right-handed: χ^2^=1.17, p=0.279; Fig. S2).

We observe that nodes with the largest number of significantly greater edges for right-handed individuals (*i.e.*, hubs) are located in both parietal lobes. Surprisingly, given the lateralization of language to the left hemisphere in the right- > left-handed group, the largest hubs are located in the secondary language regions in the right parietal lobe. In contrast, the left- > right-handed group shows hubs of significantly greater connectivity in the right hemisphere homologue of Broca’s area^36^. Overall, while the right- > left-handed group showed more widespread connectivity throughout the language networks, these edges appear to form hubs in the parietal lobe. Additionally, the cerebellum is differentially connected to the language network between groups. In particular, frontal-cerebellar connections were more prominent for the left- > right-handed group and parietal-cerebellar connections more prominent for the right- > left-handed group.

For all networks of interest, results yielded similar results when using the EHQ as a continuous or binary variable (EHQ < 0 = left-handed individuals, EHQ > 0 = right-handed individuals) (Fig. S3) and controlling for various demographic factors (e.g., age, sex) (Tables S1-S6). Overall, these results build upon previous work showing differences in activation patterns in networks of interest such that these differences are also observable in patterns of connectivity for all networks of interest.

### Generalization of networks of interest results to the PNC dataset

Using the network of interest approach, we then looked at how well these results based on data from the HBN generalized to other datasets of similar populations using data from the PNC (Fig. S4). We observed similar patterns of group differences in the HBN and PNC datasets as evidenced by the number of overlapping edges and the correlation of nodal degree between the two sets of results. First, using the hypergeometric cumulative density function to determine significance of the edge-level overlap between two networks^37^, all resulting networks of group differences were significant between the two analyses (p<0.05; Fig. S5: top row) with the exception of the network of greater edges for the left- > right-handed group in the motor network. Second, for each network of group differences, node degree--defined as the number of significant edges for each node--was calculated and correlated between the HBN and PNC results. All result pairs showed a significant correlation between nodal degree (all r’s>0.54, all p’s<0.001). Quantitatively, in the motor network, a fraction of the edges connecting the right motorstrip and ipsilateral cerebellum are present in the PNC as well. Whereas in the somatosensory network, similar crossing patterns connecting somatosensory nodes with contralateral cerebellum nodes are observed. Overlapping edges in the language network continue to highlight the importance of the cerebellum in connectivity differences between the left- > right-handed and right- > left-handed groups. Notably, the measures of handedness between the HBN and PNC were conducted differently as the HBN utilized EHQ scores which ranged from -100 to 100 while the PNC was based on self-reported measures of dominant hand for a hand tapping task. Despite differences in behavioral measures, similar patterns of connectivity were repeatedly identified as significantly different between the left- > right-handed and right- > left-handed groups. This highlights the robustness and generalizability of these results.

### Whole-brain analysis: HBN + PNC

After having established that observed differences between the left- > right-handed and right- > left-handed groups in the HBN generalize to the PNC, we combined the two datasets to increase our sample size and statistical power for whole brain analyses. To harmonize the handedness measures in the HBN and PNC, we binarized the EHQ scores to make them consistent with the PNC, such that individuals with a score below 0 were classified as primarily left-handed (left- > right-handed) and individuals with a score above 0 were classified as primarily right-handed (right- > left-handed).

Despite previous literature, widespread connectivity was observed across the whole brain between the left- > right-handed and right- > left-handed groups at the level of the whole brain (Figs. 2A & S6), beyond those in the networks of interest. Two clusters consisting of 1600 edges (right- > left-handed) and 1450 edges (left- > right-handed) exhibit significantly different (p<0.05, corrected) connectivity between the two groups. Similar to the networks of interest analyses, cerebellar connections are prominent (left- > right-handed: 39.69% significant edges, right- > left-handed: 31.78% significant edges; Fig. 2B). We observe a large proportion of edges exhibiting greater connectivity for the right- > left-handed group between the right cerebellum and the left prefrontal regions. In contrast, edges of greater connectivity for the left- > right-handed group were more localized to connections between the cerebellum and posterior regions (e.g., the occipital and parietal lobes). Results are similar when controlling for various demographic factors (e.g., age and sex for both HBn and PNC; scan sites and clinical diagnoses for HBN) (Tables. S14-S19).

**Fig. 2:**
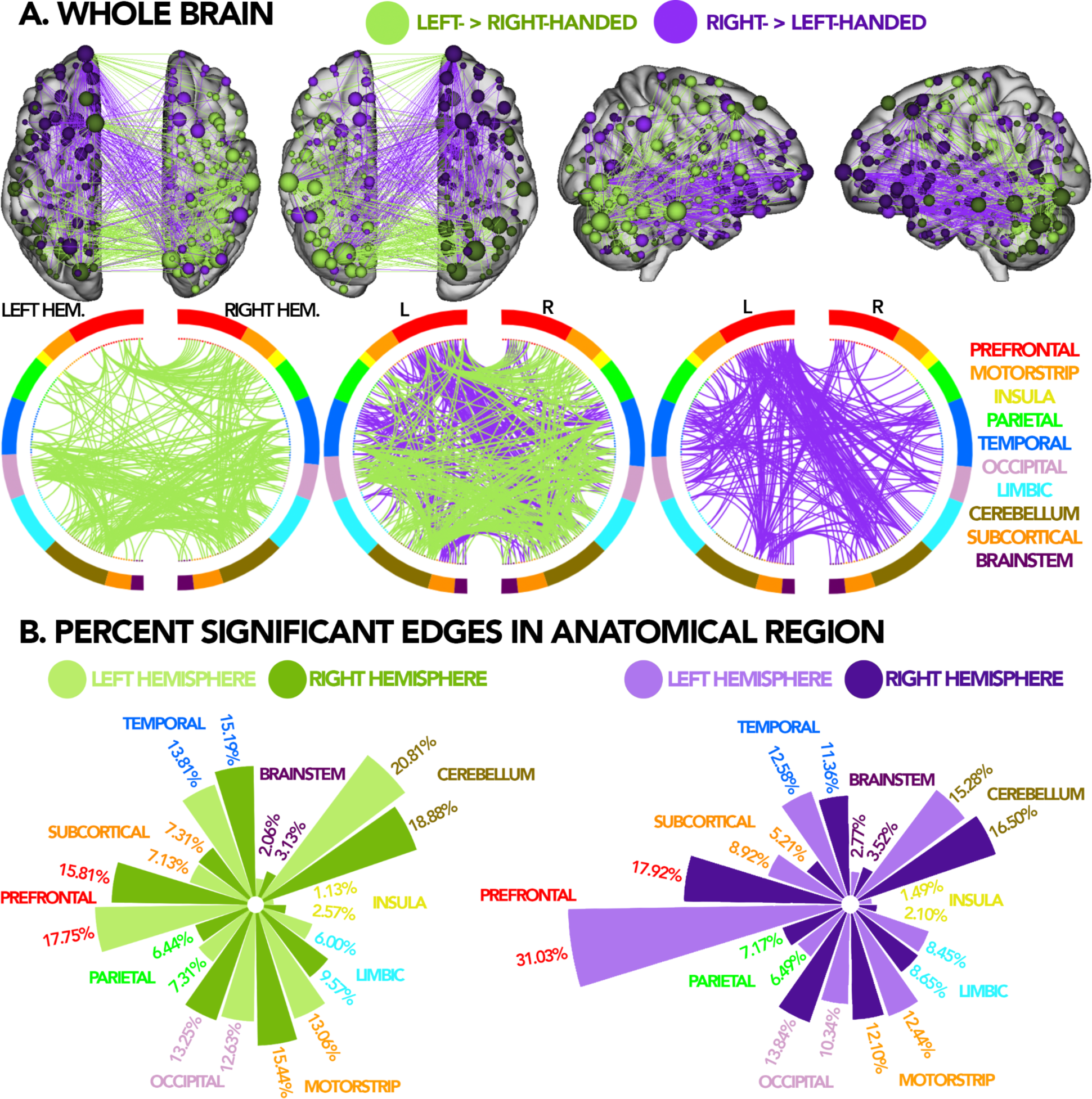
(A) Brain and circle plots for the entire connectome (p-val = 0.0018), circle plots and brain plots were thresholded at a degree threshold of 50 for visualization. Results for the left- > right-handed group are shown in green while results for the right- > left-handed group are shown in purple. Top row for each section shows significant edges drawn on an anatomical brain with nodes sized based on the number of significant edges identified. Bottom row shows circle plots where the left and right hemispheres are depicted as left and right semi-circles, respectively. The middle circle plot shows an overlay between groups. Nodes are color-coded by anatomical regions constructed based on the Shen atlas, each line depicts a significant edge identified through NBS. (B) Circular bar graph quantifying the percent of significant edges in each anatomical network corresponding with the circle plots in 2A split by left and right hemispheres, for left- > right-handed and right- > left-handed groups

Edges of greater connectivity for the right- > left-handed group were more *lateralized* within the left hemisphere (within left hemisphere: 423 edges; within right hemisphere: 324 edges;χ^2^=6.46, p=0.011; Fig. S7), consistent with the theory of left-hemisphere dominance in right-handed individuals^38^. However, edges of greater connectivity for the left- > right-handed group were not *lateralized* to either hemisphere (within left hemisphere: 353 edges; within right hemisphere: 411 edges;χ^2^=2.20, p=0.138; Fig. S7), consistent with the observation of a mix of left- and right-hemisphere dominance, or even right-hemisphere dominance, in the left- > right-handed group.

No differences in the distribution of between and within-hemisphere edges in left- or right-handed individuals are observed (left- > right-handed between: 836/1600 edges or 52.3%; within: 764/1600 edges or 47.8%; χ^2^=1.62, p=0.203; right- > left-handed between: 732/1450 edges or 49.8%; within: 737/1450 edges or 50.2%; χ^2^=0.01, p=0.920; Fig. S7).

The largest proportion of edges that differed between left- and right-handed individuals were localized to the prefrontal lobe (left- > right-handed: 33.56% significant edges, right- > left-handed: 48.93% significant edges; Fig. 2B), consistent with our network of interest results, where expressive language processing nodes (*e.g.*, Broca’s region) and secondary motor nodes are located. Surprisingly, but in line with Fig. 2A, the cerebellum contained the second largest amount of edges that differed between the left- > right-handed and right- > left-handed groups (left- > right-handed: 39.69% significant edges, right- > left-handed: 31.78% significant edges; Fig. 2B). These results were consistent when normalizing the number of edges within each network (Fig. S8). Of the 3079 edges that were identified as significantly different between the two groups at the whole-brain level, only 16.95% were also initially identified as significant using the networks of interest analysis. Overall, this observation suggests that functional connectivity differences between left- and right-handed individuals span the whole brain--rather than being localized to specific networks as suggested by previous literature^4, 6^.

Next, we quantified the effect size via Cohen’s D of the connectivity differences between left- and right-handed individuals for all edges. These effect sizes ranged from -0.3 to 0.3, consistent with the observation that brain-behavior associations tend to have low to medium effect sizes^39, 40^. To help put these whole-brain differences into comparable context relative to sex differences, for primarily right-handed individuals only, we compared the connectomes between male and females participants (based on self-reported sex) and quantified edgewise effects sizes for these differences. Broadly, the effect sizes observed for sex differences in whole-brain functional connectivity were of a similar magnitude as the effect sizes observed for handedness differences (Fig. 3) with no significant differences between the two distributions of effect-sizes being observed. Together, these results suggest that handedness differences account for a similar amount of individual differences in the connectome as sex differences, and underscore that the handedness effects are neurobiogically meaningful in addition to being statistically significant.

**Fig. 3:**
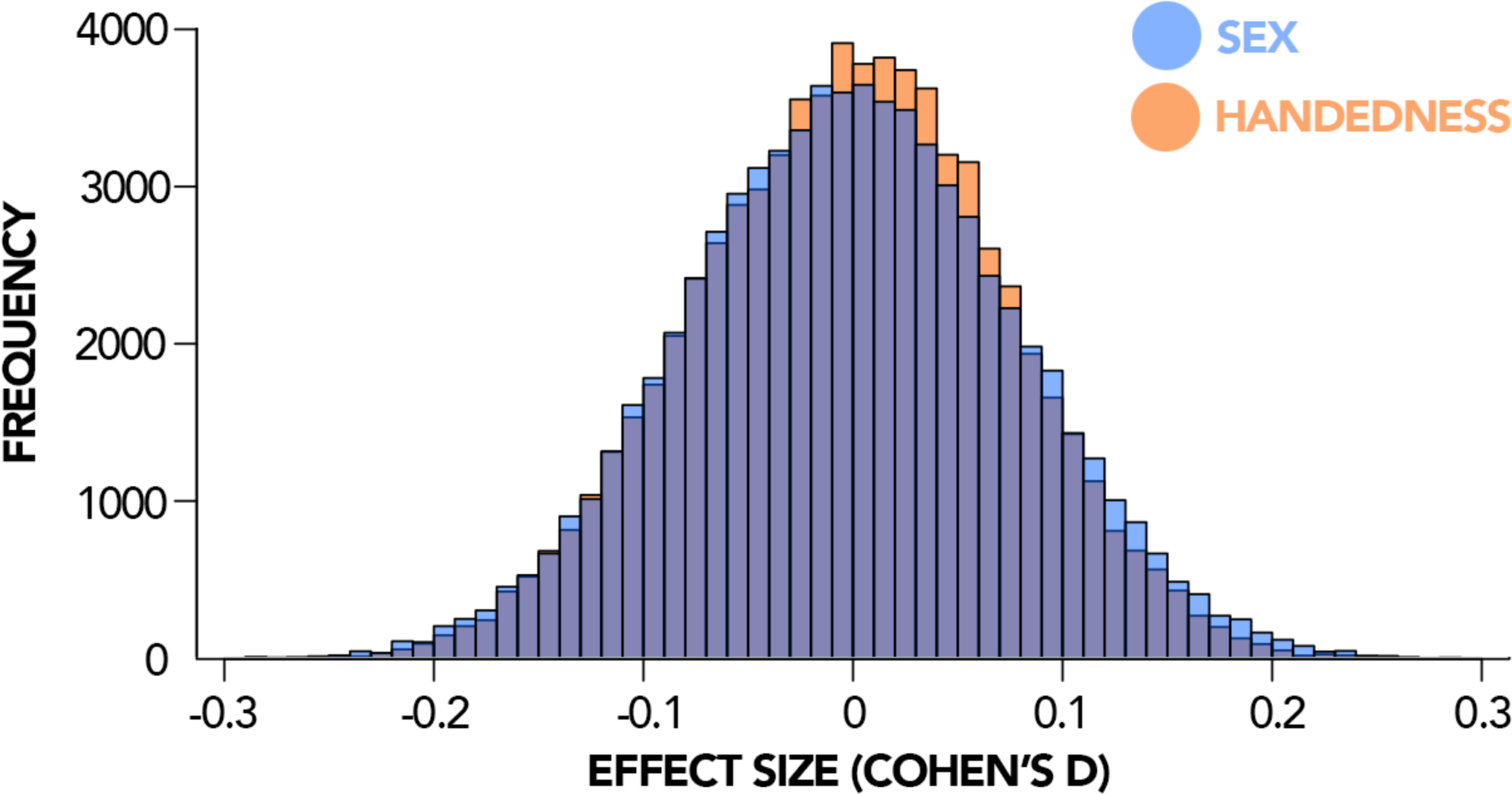
Comparison of effect sizes for each edge in a connectome (total 35,778 edges) plotted onto a histogram for handedness and sex.

### Between and within hemispheres

Analyses on between-hemisphere edges (Fig. 4A) demonstrate very similar patterns to those of the whole brain. The same bundles of edges forming between the left prefrontal and contralateral cerebellum make up the majority of between-hemisphere edges that are significantly greater for the right- > left-handed group. Similarly, the same patterns of cerebellar edges for the left- > right-handed group is observed in our between-hemisphere analyses.

**Fig. 4:**
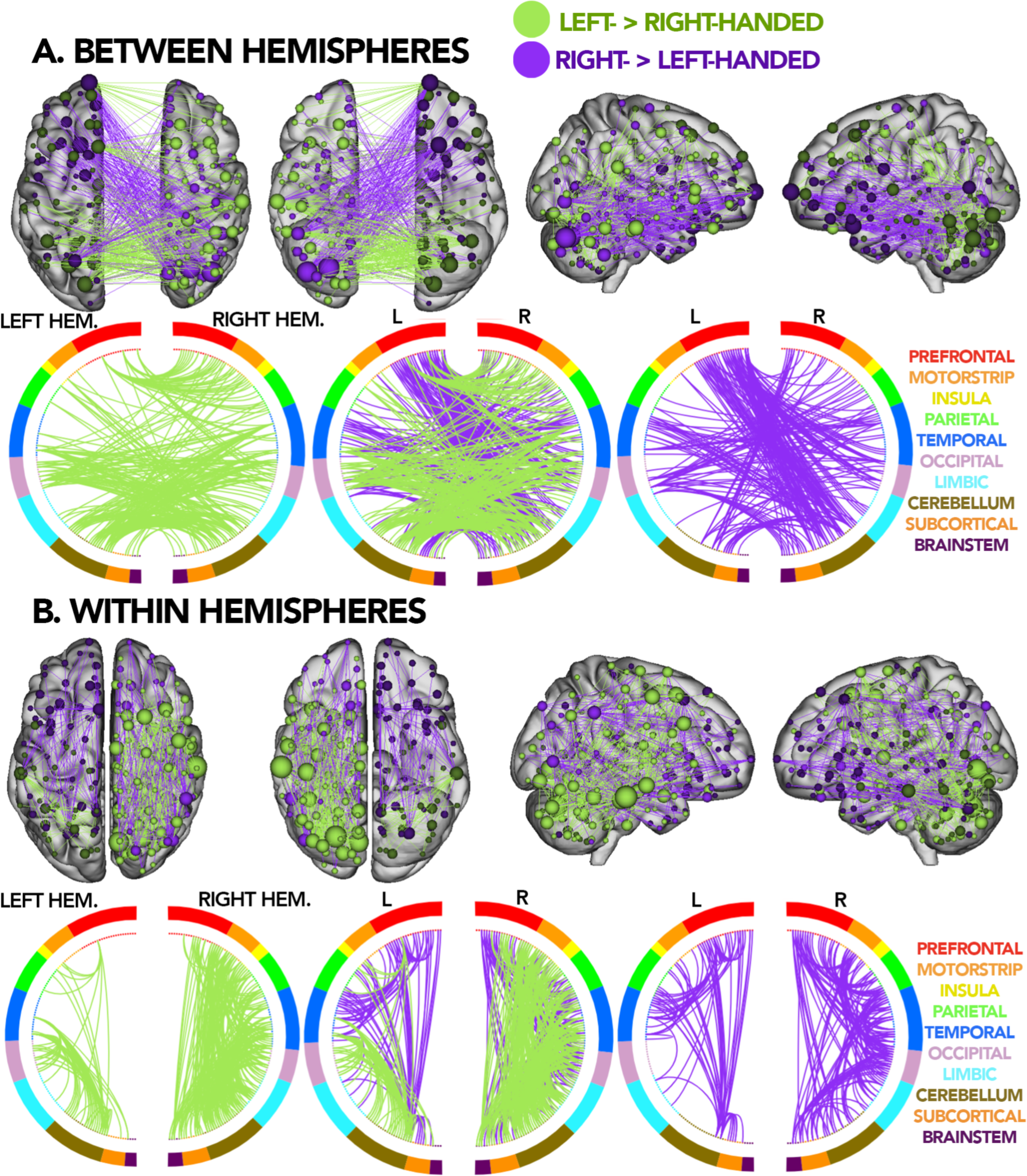
(A) Brain and circle plots for significant edges between-hemispheres (p = 0.012), circle plots and brain plots were thresholded at a degree threshold of 25 for visualization. (B) Brain and circle plots for significant edges for within-hemispheres (left hemisphere p = 0.019, right hemisphere p = 0.022), circle plots and brain plots were thresholded at a degree threshold of 25 for visualization. Results for the left- > right-handed group are shown in green while results for the right- > left-handed group are shown in purple. Top row for each section shows significant edges drawn on an anatomical brain with nodes sized based on the number of significant edges identified. Bottom row for each section shows circle plots where the left and right hemispheres are depicted as left and right semi-circles, respectively. The middle circle plot shows an overlay of circle plots for both groups. Nodes are color-coded by data- driven networks constructed based on the Shen atlas, each line depicts a significant edge identified through NBS.

Overall, cerebellar edges make up the majority of significant between-hemisphere edges when comparing the two groups (1108/1568 edges or 70.7%).

The within-hemisphere results show diverging *laterality* patterns from our whole-brain analyses (Fig. 4B). Consistent with the whole-brain results, edges of greater connectivity for the right- > left-handed group were more *lateralized* within the left hemisphere (within left hemisphere: 423 edges; within right hemisphere: edges: 324;χ^2^=6.46, p=0.01). However, edges of greater connectivity for the left- > right-handed group were more *lateralized* within the right hemisphere for within-hemisphere edges (within left hemisphere: 324 edges; within right hemisphere: 821 edges;χ^2^=97.4, p<0.001). This finding is in contrast to our whole-brain results, where the left- > right-handed group exhibited a non-significant lateralization to the right hemisphere. Perhaps, given the more localized analysis to only within-hemisphere, the null clusters from the permutation analysis were smaller, leading to additional information surviving NBS correction. In other words, by restricting our analysis, we were able to see better under the spotlight^43^.

Together, these results are consistent with the observation of left-hemisphere dominance in primarily right-handed individuals and mixed or right-hemisphere dominance in primarily left- handed individuals^4, 44^.

### Cerebellum

Given the striking contribution of the cerebellum to the network of interest (Fig. 1) and whole-brain group differences (Figs. 2 & 4) as well as the relatively unexplored functional differences in the cerebellum between the left- > right-handed and right- > left-handed groups^45^, we further investigated cerebellar differences in functional connectivity using our networks of interest approach.

Within the cerebellar network (Fig. 5), clusters consisting of 463 edges (right- > left-handed) and 558 edges (left- > right-handed) exhibit significantly different (p<0.05, corrected) connectivity between the left- > right-handed and right- > left-handed groups. The left- > right-handed group show large bundles of edges with significantly greater connectivity between the cerebellum and the motor strip and somatosensory areas, consistent with results for the motor and language networks (Fig. 1). Interestingly, edges of greater connectivity for the left- > right-handed group are generally confined towards the posterior regions of the brain whereas edges of greater connectivity for the right- > left-handed group are generally confined to the frontal regions.

**Fig. 5:**
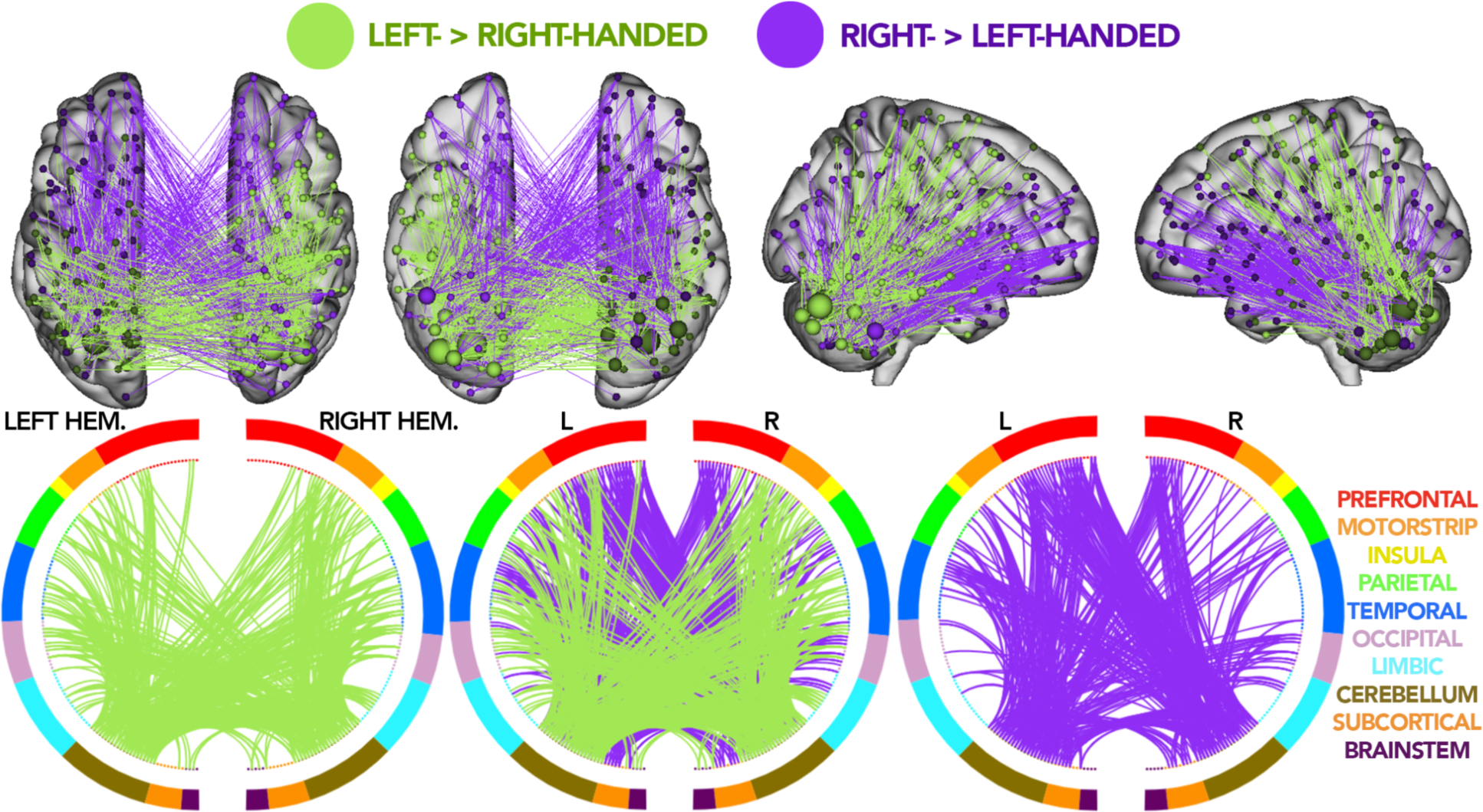
Brain and circle plots for all nodes in the cerebellum (p-val = 0.031). Results for the left- > right-handed group are shown in green while results for the right- > left-handed group are shown in purple. Top row shows significant edges drawn on an anatomical brain with nodes sized based on the number of significant edges identified. Bottom row shows circle plots where the left and right hemispheres are depicted as left and right semi-circles, respectively. The middle circle plot shows an overlay between left- and right-handed individuals. Nodes are color-coded by data- driven networks constructed based on the Shen atlas, each line depicts a significant edge identified through NBS.

Results are similar when controlling for various demographic factors (e.g., age, sex) (Tables. S1-S4). Edges of greater connectivity in the right- > left-handed group are mostly between-hemisphere edges rather than within hemispheres (between: 274/463 edges or 59.2%; within: 189/463 edges or 40.8%; χ^2^=7.69, p=0.006; Fig. S12) and are more *lateralized* within the left hemisphere (within left hemisphere: 126 edges; within right hemisphere: 63 edges;χ^2^=14.86, p<0.001; Fig. S12). No differences in the distribution of edges of greater connectivity in the left- > right-handed group were observed (between: 284/558 edges or 50.9%; within: 274/558 edges or 49.1%; χ^2^=0.09, p=0.76; within left hemisphere: 132 edges; within right hemisphere: 152 edges;χ^2^=0.71, p=0.40; Fig. S12).

## Discussion

Using functional connectomes from two large open-source datasets (the Healthy Brain Network and Philadelphia Neurodevelopmental Cohort), we show that differences in the functional organization between groups of primarily left- and primarily right-handed individuals are found not only in previously identified functional networks, but in every brain region with a strikingly large amount of differences for edges incident to the cerebellum. We began by investigating differences in networks of interest, as established by previous activation studies, to show that these differences can also be detected by functional connectivity. These differences also robustly generalized across datasets. In a combined sample from both datasets, we show that differences in functional connectivity between the left- > right-handed and right- > left-handed groups are present across the whole brain. In particular, to emphasize the significance of these differences, we compared the distribution of effect sizes to those from self-reported sex.

Handedness differences exhibit similar effect sizes as sex differences suggesting handedness may be a factor researchers should control for in future large-scale connectome studies. Finally, while previous studies have focused on the cortex^28, 46^, we find that the most striking differences between the left- > right-handed and right- > left-handed groups are edges located within and between the cerebellum. Together, these results characterize fundamental differences in the functional connectome associated with handedness.

### Whole brain analyses: going beyond regions of interest

Deviating from traditional region and network of interest approaches, our whole-brain results emphasize that differences between the left- > right-handed and right- > left-handed groups are wide-spread across the whole brain rather than localized to a few regions and networks. Indeed, only 16.95% of edges from the whole-brain results were identified as significant using the networks of interest analysis. The widespread nature of our results is also in contrast to emerging morphometric studies of handedness, which similarly report sparse, localized differences between the two groups^47^. A potential explanation may be that functional connections have greater neuroplasticity than anatomical structures^34^. Given the relative rarity of left-handed individuals (approximately 10% of the population^48^), they may be forced to use tools designed for right-handed individuals (e.g., scissors or computer mouse). This adaptation likely results in neuroplasticity with large-scale changes in functional connectivity, likely not observable in fixed anatomical structures.

### Cerebellum

Despite a majority of handedness work focusing on the cortex^2,28, 32, 49–52^, the cerebellum demonstrated the second largest number of significant edges of networks evaluated in the data-driven, whole-brain analysis (the prefrontal lobe, which includes several of our networks of interest, contained the largest number of significant edges). Reported associations between handedness and the cerebellum are limited^13, 45^. Perhaps this result is not surprising given the cerebellum’s role in motor control^53, 54^ and the association of motor control and handedness^28, 49^. Nevertheless, most of the significant edges do not involve the motor cortex, in line with the recent trend to consider the cerebellum as a cognitive region, rather than a solely motor region^55^. The cerebellum develops rapidly postnatally^56^, during a time when infants acquire a vast amount of skills and handedness begins to crystalize. As periods of rapid development show the greatest neuroplasticity^34^, it may be reasonable to expect that functional connections are more plastic in the cerebellum than other regions, resulting in the large functional differences in the cerebellum.

### Effect sizes: controlling for handedness in large studies

Given the magnitude of effect sizes in neuroimaging and clinical and social factors associated with sex differences^57^, sex is routinely controlled for in neuroimaging studies^58, 59^. The similarity between the effect size magnitude of handedness differences and sex differences in FC underscores the importance of potentially accounting these functional differences. Future studies that include a large number of left-handed individuals may need to control for handedness in a similar manner as other covariates, such as sex. One caveat might be that left-handed individuals are relatively rare (around 10% of the population^1^). Many functional connectivity studies may not have a sufficiently large sample of left-handed individuals to properly estimate these effects. However, potential differences in the connectome should not be used to justify excluding left-handed individuals from a study. Best practices in maintaining representative samples necessitates the inclusion of left-handed individuals^60^. Nevertheless, the best approach for accounting for handedness differences in the connectome remains to be determined.

### Functional connectivity relative to other brain studies of handedness

In line with previous results from activation, morphometric, neuropsychological, and lesion^61^ studies, we found that functional connectivity incident to the motor, somatosensory, and language networks differed between primarily left- and primarily right-handed individuals. While our results build upon this previous work, differences in functional connectivity do not necessarily translate to observed differences in brain activation^62^ or structure. For instance, one may expect large functional connectivity differences in Broca’s and Wernicke’s areas^3, 32^ based on previous work in activation studies regarding lateralization differences in language between left- and right-handed individuals. Yet, we found the largest number of significantly different edges clustered in the right-hemisphere, located in secondary language processing regions of the temporoparietal junction (in the right- > left-handed group). The lack of one-to-one translation of results between functional connectivity and activation likely holds in the other direction, too. In other words, the lack of differences in functional connectivity does not imply that activation patterns in Broca’s or Wernicke’s areas between the left- > right-handed and right- > left-handed groups during a language task would be the same. Patterns of within- and between-hemisphere edges also appear to be consistent across all three networks.

### Lateralization/cross hemispheric connections

Additionally, while little lateralization was observed using the network of interest, strong lateralization effects were observed in the whole-brain results, consistent with patterns of left hemisphere dominance in right-handed individuals^44, 63^. As such, we delved deeper into analyzing patterns of connectivity for significant edges within and between hemispheres. We consistently observe a greater amount of within-hemisphere edges for the left- > right-handed group whereas we observe a greater amount of between-hemisphere edges for the right- > left-handed group. These differences in between and within-hemisphere edges could result from differences in corpus callosum connectivity associated with handedness. Differences in the amount of between-hemispheric connections via the corpus callosum has been shown to be linked to the extent of handedness an individual exhibits^7^ (i.e., the more ambidextrous an individual, the more connections between hemispheres). Overall, these observations highlight that handedness differences in the functional connectome are vastly more distributed than previously understood.

### Strengths and weaknesses

There are several notable strengths of our study. First, we used two large open-source datasets, allowing for a large sample of left-handed individuals (n>225), the application of whole-brain approaches and the ability to investigate generalization/replication of results across study designs. Without the large sample size and whole-brain analyses, important results (e.g., the widespread nature of handedness differences and the large handedness differences in cerebellum) may not have been discovered. Similarly, generalizing results from the HBN to the PNC highlight their robustness, especially considering the different handedness measures across the datasets. Second, by focusing on school-age children and adolescents as opposed to adults, we can better investigate the innate differences in connectivity, rather than adaptive differences acquired over the course of life. For example, historically, left-handed individuals were often forced to write right-handed. Also cultural sigma may have led others to become functionally right-handed^64^ (e.g., sinister means both evil and left). Yet, even in a younger sample, fully ruling out adaptive differences is not possible.

Nevertheless, there are several notable limitations of our study. First, while all of our analyses are based on the same procedure and thresholds using NBS, it is important to note that running NBS on a subsetted connectome as opposed to the whole connectome will select different edges as a result. For instance, an edge that is initially identified as significant based on a subsetted connectome (like in our networks of interest) may not be identified as significant when using the entire connectome. Second, in defining our networks of interest, we based our definitions on differences in activation patterns shown in previous studies^6, 8, 65^. These previous studies have typically reported their results in the context of Brodmann areas, where our connectomes are parcellated based on a 268-node functionally defined atlas. Thus, we manually identified nodes that overlapped with these Brodmann areas, however, due to the differences in the Shen atlas and the Brodmann areas, our networks of interest may not have captured the exact regions that were reported in previous studies. Moreover, the Brodmann atlas is symmetrical between hemispheres, while our 268-node atlas is not. As such, there are also asymmetries between areas of the brain included in our analyses of networks of interest between the left and right hemispheres. Third, in our whole-brain analysis, we were limited to binarizing the EHQ in the HBN datasets for harmonization with the PNC handedness measure. While we could have explored a third group of ambidextrous individuals in HBN, we were limited by: (a) the fact that there is no gold standard for the range of scores in the EHQ to classify an ambidextrous group^66^ and (b) the PNC’s measures of handedness was a forced-choice self report of handedness. Because of the variability and range in EHQ scores, we chose to conduct our initial analyses on the HBN and subsequent generalization/harmonization to the PNC. Finally, to address handedness interactions with sex and age, we repeated all NBS analyses using partial correlation to control for these factors (sex: Table S1, age: Table S3) as well conducting combined analyses for the two datasets separately (sex: Table S2, age Table S4). These results robustly demonstrate that while sex and age are potential confounding factors, our results remain unchanged as the same significant edges are identified with and without controlling for these factors. Additionally, we also controlled for scanning site and clinical diagnoses for the HBN, since this population was scanned across multiple sites and contained many subjects with clinical diagnoses (site: Table S5, diagnoses: Table S6). Similarly, the same significant edges were robustly identified as significant with or without controlling for scanning site and clinical diagnoses.

### Future directions

In sum, we show that differences in the functional connectome associated with handedness are distributed across the brain, including previously unreported differences associated with the cerebellar network. Future directions include investigations into sex-handedness interaction^11, 28, 67, 68^ (as majority of left-handed population consists of males^48^), into a third ambidextrous group, and into potential interactions between handedness and psychiatric diagnoses (as non-right handedness is overrepresented in various psychiatric disorders, namely schizophrenia^25^). As the observed differences show meaningful effect sizes, future studies may need to consider accounting for handedness. This work serves as a starting point to account for handedness in functional connectivity studies, in particular for studies involving neuropsychiatric disorders.

## Methods

### Dataset: HBN

All connectomes for initial analyses (Fig. 1) were generated from resting-state scans obtained from the Healthy Brain Network (HBN)^23^. All resting-state scans are 10 mins long using a 1.5 T Siemens Avanto system equipped with 45 mT/m gradients in a mobile trailer at four different sites around the New York greater metropolitan area: Staten Island, Cornell University, City University of New York, and Rutgers University. After excluding subjects for missing scans/data and excessive motion (>0.2 mm), 905 subjects remain (right- > left-handed group: 787, left- > right-handed individuals: 118). Subjects’ ages ranged from 5-22 where 111 subjects had no diagnosis and 794 had some diagnosis of learning disorders or symptoms of psychiatry.

Edinburgh Handedness Questionnaire scores were used as a measure of the extent subjects were left-handed and right-handed. Scores ranged from -100 to 100 where -100 is considered an extremely left-handed individual and 100 is considered an extremely right-handed individual.

### Dataset: PNC

For generalization, we used data from the Philadelphia Neurodevelopmental Cohort (PNC)^24, 26^ by following the same preprocessing pipelines used with HBN. All resting-state scans are 6 mins long using a single 3T Siemens TIM Trio whole-body scanner with the VB17 revision of the Siemens software. All participants were scanned at the University of Pennsylvania in Philadelphia, PA. After excluding subjects for missing scans/data and excessive motion (>0.2 mm), 859 subjects remain (right- > left-handed individuals: 742, left- > right-handed individuals: 117). Subjects’ ages ranged from 8-23 yrs and measures of handedness were based on self-reports of dominant hand to complete another finger tapping task in the dataset (data not used in our analyses).

### Preprocessing and generating connectomes

Both the HBN and PNC datasets were analyzed with identical processing pipelines. Structural scans were first skull stripped using an optimized version of the FMRIB’s Software Library (FSL)^69^ pipeline^70^. Functional images were motion corrected using SPM12. All further analyses were performed using BioImage Suite^71^ and included linear and nonlinear registration to the MNI template, unless otherwise specified. Several covariates of no interest were regressed from the data including linear and quadratic drifts, mean cerebral-spinal-fluid (CSF) signal, mean white-matter signal, and mean gray matter signal. For additional control of possible motion-related confounds, a 24-parameter motion model (including six rigid-body motion parameters, six temporal derivatives, and these terms squared) was regressed from the data. The data were temporally smoothed with a Gaussian filter (approximate cutoff frequency=0.12 Hz).

Nodes were defined using the Shen 268-node brain atlas^72^, which includes the cortex, subcortex, and cerebellum as described in prior CPM work. The atlas was warped from MNI space into single-subject space via a series of linear and non-linear transformations. Resting state connectivity was calculated on the basis of the ‘raw’ task time courses^73^, which emphasizes individual differences in connectivity^43^. This involved computation of the mean time courses for each of the 268 nodes (i.e., averaging the time courses of all constituent voxels).

Node-by-node pairwise correlations were computed, and Pearson correlation coefficients were Fisher z-transformed to yield symmetric 268x268 connectivity matrices, in which each element of the matrix represents the connectivity strength between two individual nodes (i.e., ‘edge’).

### Defining Networks of Interest

Based on previous literature on differences in handedness^6, 8, 27–32, 49, 74^, we defined three networks of interest: motor, somatosensory, and language using the Brodmann Areas that were reported for each publication (Table 1). Connectomes were partitioned into matrices that only contained edges that stem from a node of interest or edges between nodes of interest.

Differences between the left- > right-handed and right- > left-handed groups were estimated using Network-Based Statistics^17^ (component-determining threshold z=1.96, 2-tailed, K=5000 permutations) for each network separately.

Initial analyses conducted on the HBN utilized raw EHQ scores to identify differences between the left- > right-handed and right- > left-handed groups (Fig. 1), whereas analyses done purely on the PNC relied on 0/1 self-reported measures of handedness to replicate the same analyses (Fig. S3).

### Generalization to the PNC

The significance of the overlap between the networks of interest and the whole brain between the HBN and PNC was determined with the hypergeometric cumulative density function^37^, which returns the probability of drawing up to x of K possible items in n drawings without replacement from an M-item population. This was implemented in Matlab as: p=1-hygecdf(x, M, K, n), where x equals the number of overlapping edges, K equals the number of connections in the HBN network of interest, n equals the number of connections in the PNC network of interest, and M equals the total number of edges in the matrix (35,778). Percent overlap for the barplots (Figs. S5 and S9) were calculated as: number of overlapping edges/(HBN significant edges + PNC significant edges - overlapping edges).

### Combined analyses: HBN + PNC

For all analyses where we combined data from the HBN and PNC, we addressed incongruencies in handedness measures by binarizing EHQ scores such that subjects who scored below 0 were considered primarily left-handed and above 0 were considered primarily right-handed. No subject had an EHQ score of exactly 0. Networks of interest analyses conducted on thresholded HBN EHQ scores at 0 (Fig. S4) exhibited similar patterns of connectivity both in the brain and circle plots as analyses conducted on raw EHQ scores (Fig. 1).

Whole brain functional connectivity differences between left- and right-handed individuals were estimated using the Network-Based Statistic^17^ (component size statistic; component-determining threshold z = 1.96, 2-tailed, K=5000 permutations) for each network separately.

Due to the large number of edges identified as significant in our whole-brain analyses, we thresholded our visualizations for the brain and circle plots (Fig. 2A) to degree threshold 50, brain plots with varying thresholds are shown in SI (Fig. S6). The remainder of our whole-brain analyses showing the number of significant edges in each anatomical region is based on the full connectome without thresholds.

Similarly, our between and within-hemisphere brain and circle plots (Fig. 4) were thresholded at degree threshold 25 to demonstrate patterns of connectivity. Visualizations with varying thresholds are shown in SI (Fig. S10). All quantifications are based on subsetted connectomes to include only between or within-hemisphere edges without thresholds.

Finally, our analyses on the cerebellum brain and circle plots (Fig. 5) show the full set of significant edges without thresholds to demonstrate patterns of connectivity. All quantifications are based on subsetted connectomes to include edges between and within the cerebellar nodes.

### Effect Size Comparisons

In comparing effect sizes between sex and handedness (Fig. 3), effect sizes were calculated for each edge in a 268x268 connectome across all subjects in both datasets for sex and for handedness.

### Controlling for confounding factors in datasets

Because we used developmental datasets with some clinical diagnoses to study a normative trait. We conducted additional analyses using NBS partial correlations to control for sex (Tables S1, S2) and age (Tables S3, S4) in both the HBN and the PNC to demonstrate the same significant edges were identified between the left- > right-handed and right- > left-handed groups while controlling for these differences. Results were split up into two tables when controlling for sex and age to demonstrate overlaps when analyses were conducted identically to results section of the paper (sex: Table S1, age: Table S3) and to show results still hold up when we run NBS on the two datasets, HBN and PNC, for whole brain and cerebellum (sex: Table S2, age: Table S4) separately. We also controlled for scan site (Table S5) and clinical diagnoses (Table S6) in the HBN since this sample was collected from many different scan sites and the population was biased towards clinical diagnoses (794/905 subjects had at least one clinical diagnosis). Pearson correlations were calculated between matrices of significant edges identified by NBS alone and NBS correlation controlling for each factor. Unlike our generalizations from HBN to the PNC, we opted to use correlations as opposed to hypergeometric cumulative density function (as previously used to show generalization across datasets) because samples were not independent of each other.

## Supplementary Results

### Generalization of whole-brain results between the HBN and PNC

To verify that results generalized across datasets at the level of the whole brain, rather than just within the networks of interest, we performed cluster-based inference on the entire connectome for the HBN and PNC separately, then proceeded to look at the number of overlapping edges, in the same manner as above. Generalization between the two datasets revealed significance across the whole brain and across all data-driven functional networks, analogous to functional networks of interest (Fig. S9). Across all canonical brain networks^41, 42^, the number of overlapping edges far exceeds the minimum level required for significance, with the exception of the salience network in the right- > left-handed group. At the whole brain level, slightly greater than 3% of the total number of edges overlap between the HBN and PNC results. Given that a connectome has 35,778 unique edges, it is exceedingly rare to choose a single edge out of two random draws from all edges, let alone the 91 overlapping edges we observe. Together with the network of interest results, these results repeatedly demonstrate that edges observed to significantly differ between the left- > right-handed and right- > left-handed groups are highly generalizable across datasets, despite differences in study design including, handedness measure and scanner/scan sites.<u>Between and within hemispheres</u>

A common observation from our whole-brain analyses was that the left- > right-handed and right- > left-handed groups differed in the patterns of edges forming between and within- hemispheres. Thus, we performed NBS on connectomes subsetted for within- and between networks separately.

### Supplementary Figures

**Fig. S1:**
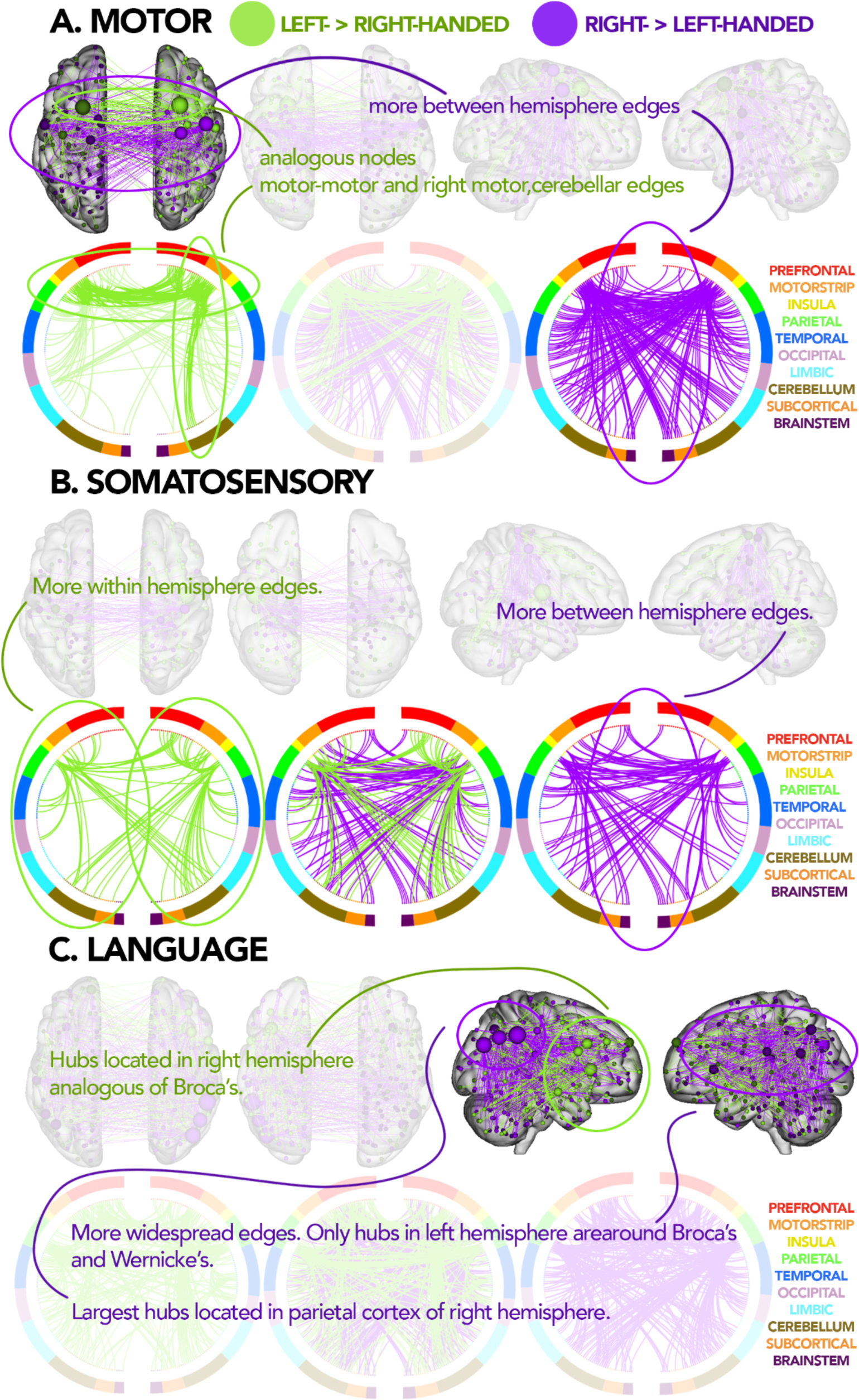
Summarization of results from Fig. 1 showing only patterns of interest.

**Fig. S2:**
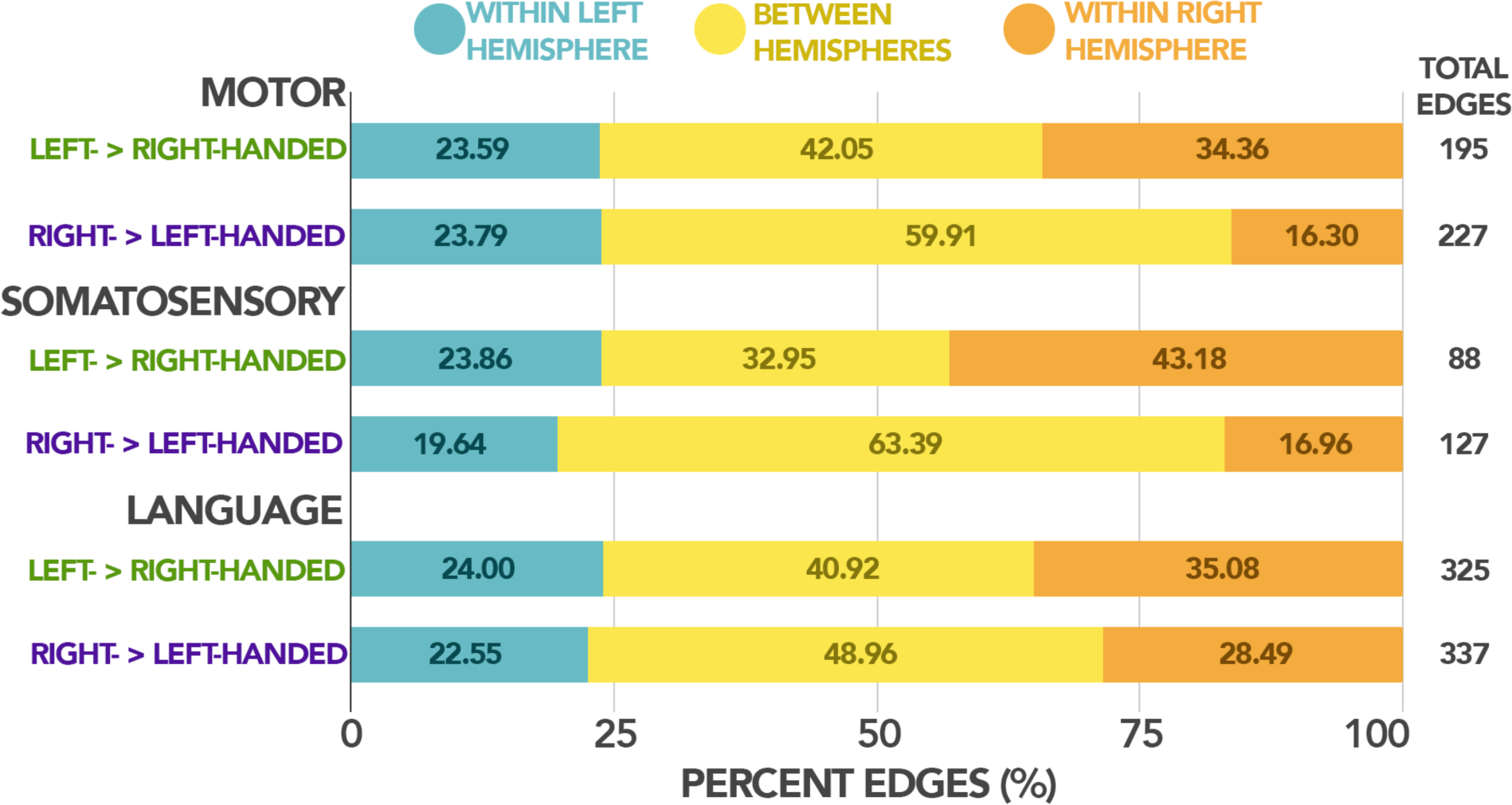
Percent of edges in each network of interest: motor, somatosensory, and language, split by left- > right-handed group and right- > left-handed group connecting within left/right hemispheres or between-hemispheres.

**Fig. S3:**
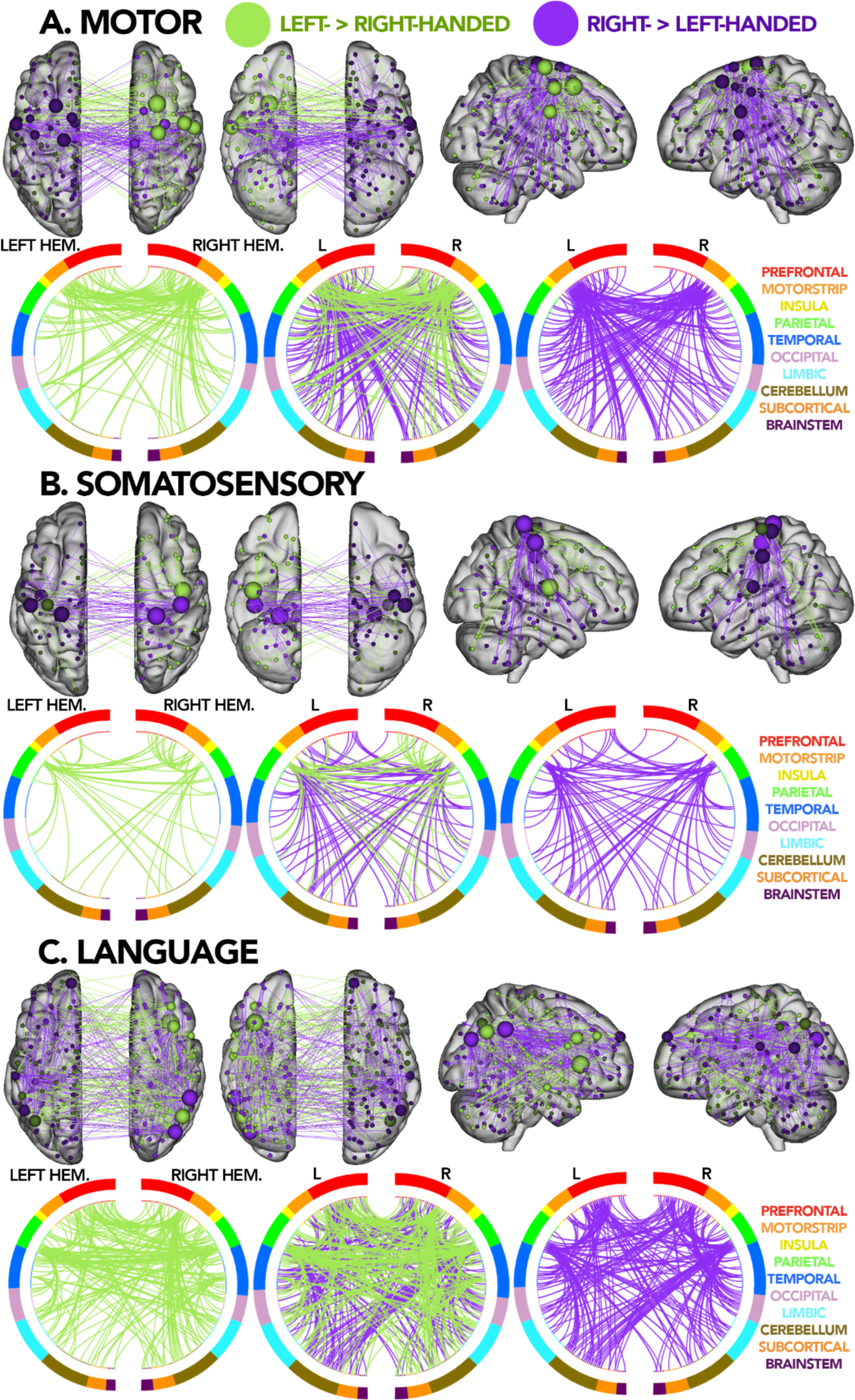
Replication of same networks of interest conducted in Fig. 1 but with raw EHQ scores converted to 0/1, thresholded at 0. EHQ scores < 0 = left- > right-handed group, EHQ scores > 0 = right- > left-handed group.

**Fig. S4:**
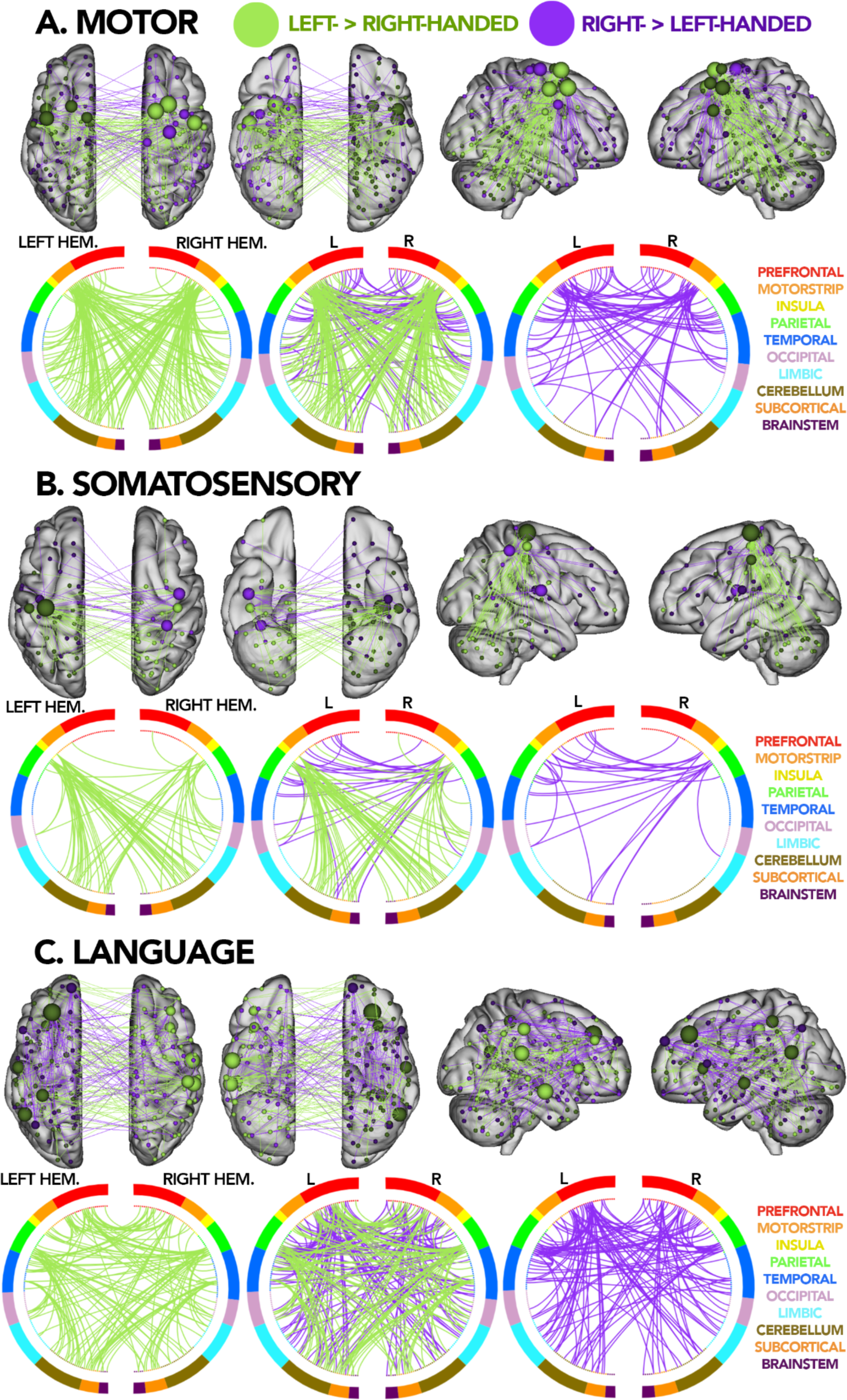
Replication of same networks of interest conducted in Fig. 1 but with data from the PNC. Behavioral scores were binarized for left- > right-handed and right- > left-handed participants.

**Fig. S5:**
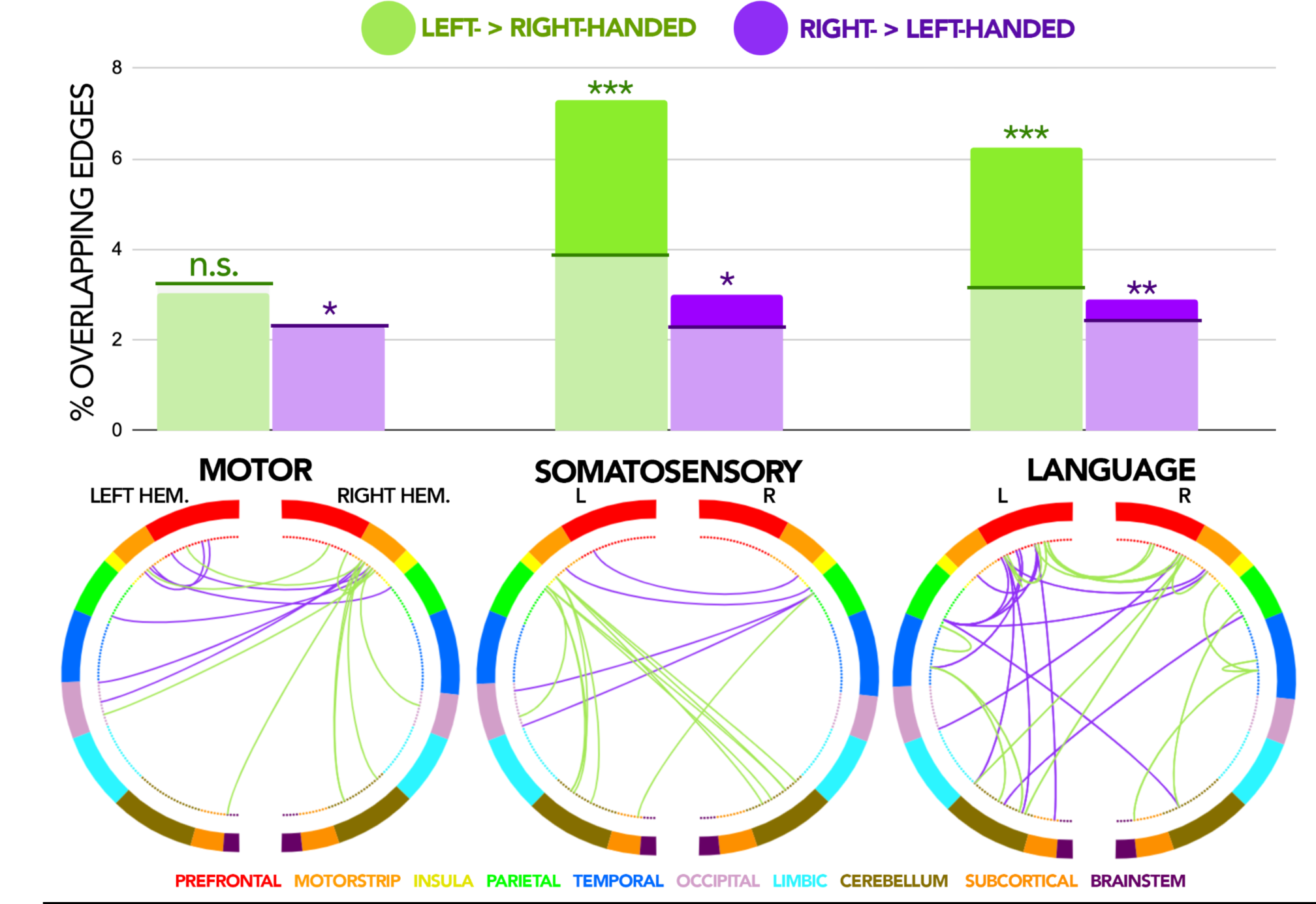
Bottom row shows circle plots where the left and right hemispheres are depicted as left and right semi- circles, respectively. Nodes are color-coded by anatomical networks constructed based on the Shen atlas, each line depicts a significant edge identified through NBS. Legend for which anatomical region each color represents is shown in a line above the three circle plots. Each circle plot shows significant edges that are present in both HBN and PNC for each network of interest: motor, somatosensory, and language, respectively. Top row depicts a bar graph of % overlapping edges for each network of interest. Lines and shaded regions in each bar indicate the minimum % of overlapping edges required for significance. * on top of each bar indicates significance where n.s. indicates not significant, *indicates p ≤ 0.05, **indicates p ≤ 0.01, and ***indicates p ≤ 0.001

**Fig. S6:**
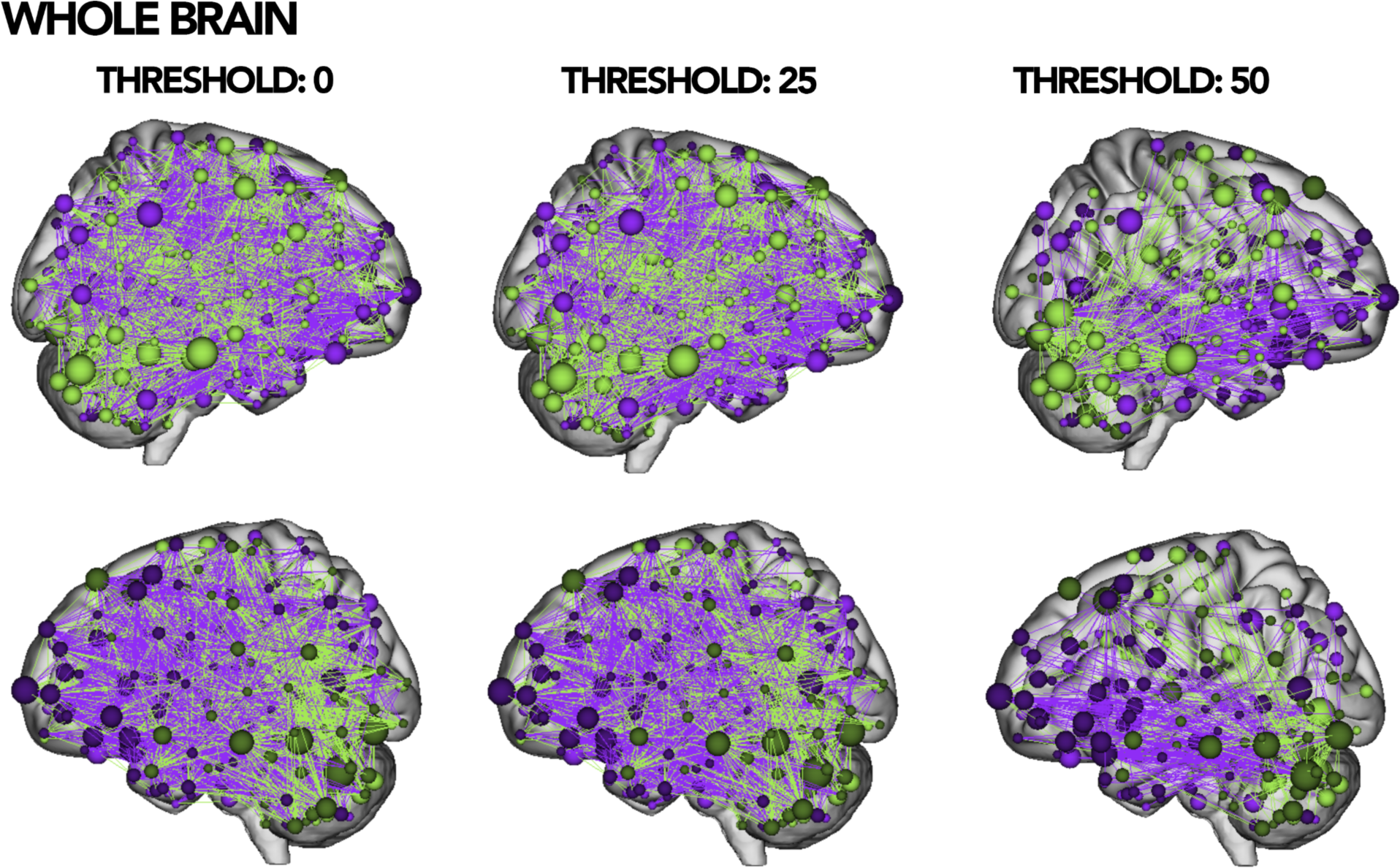
Ball-stick brain plots for differences between left-handed individuals and right-handed individuals at the level of the whole brain for the combined analyses with HBN and PNC thresholded at degree = 0, 25, and 50.

**Fig. S7:**
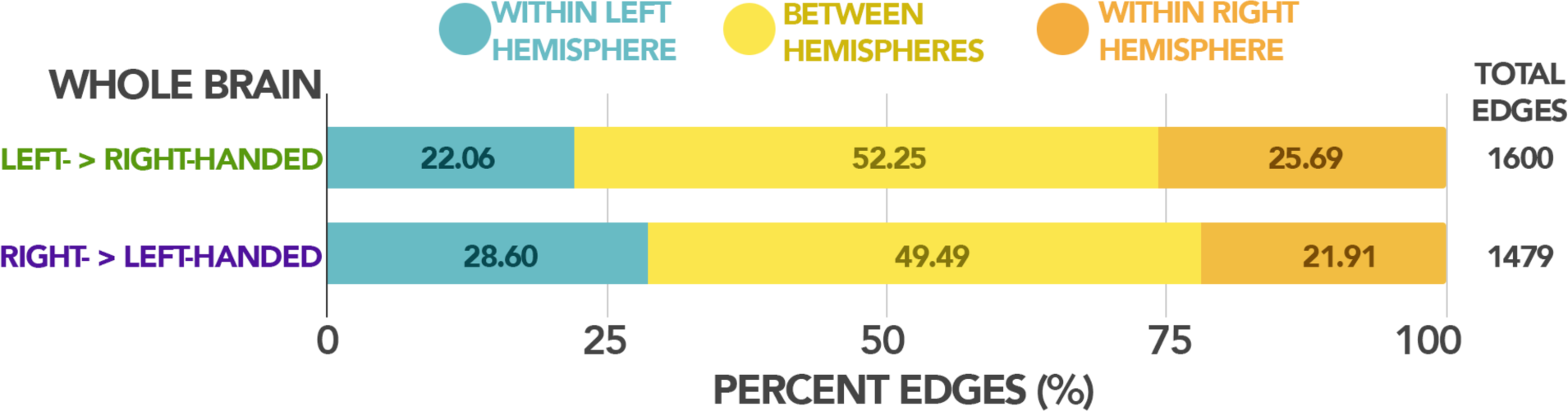
Percent of edges for left- > right-handed and right- > left-handed group for edges connecting within left/right hemispheres or between-hemispheres.

**Fig. S8:**
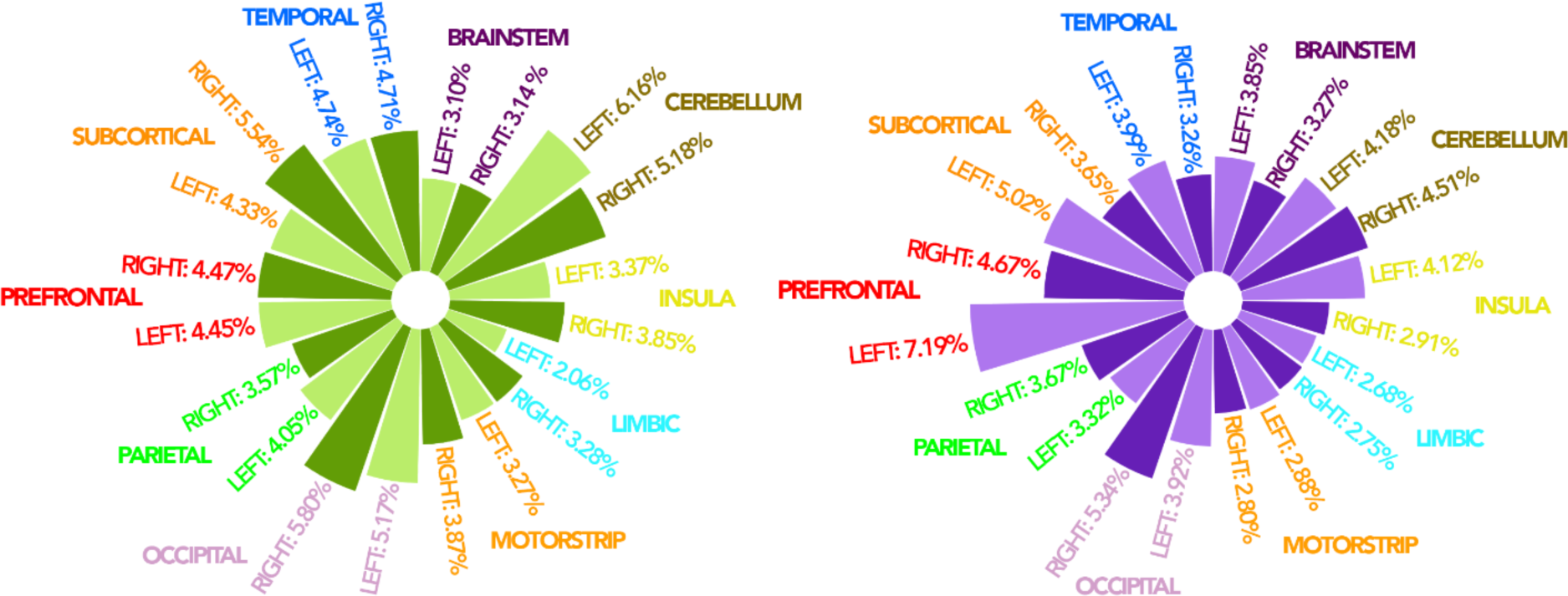
Percent of edges that identified as significant in whole brain analysis out of the total number of edges in each anatomical region, split by left and right hemispheres. Results for left-handed individuals shown in green circular bar graph on the left and results for right-handed individuals shown in purple circular bar graph on the right. Left hemispheres shown in lighter colored bars, right hemispheres shown in darker colored bars.

**Fig. S9:**
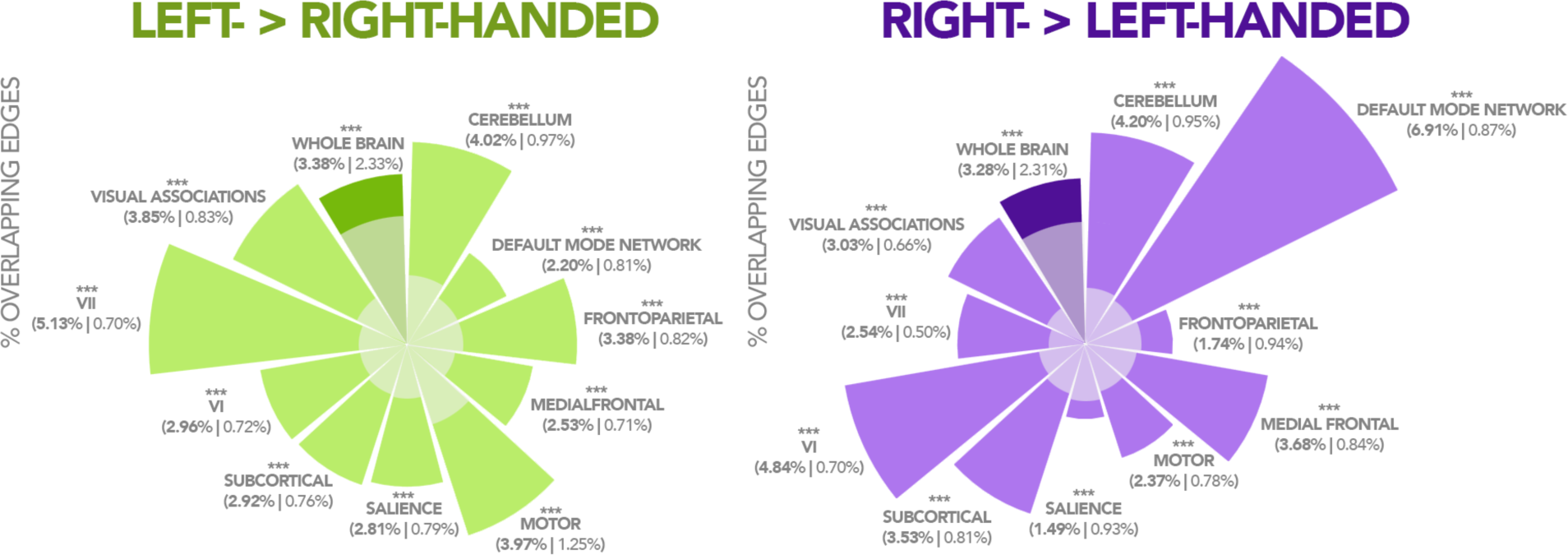
Circular bar graphs showing overlapping edges between HBN and PNC by canonical data-driven networks: Medial Frontal, Frontoparietal, Default Mode Network, Motor, VI, VII, Visual Association, Salience, Subcortical, and Cerebellum. The whole brain is shown in a darker colored bar for both the left- > right-handed and right- > left-handed group. Exact percentage of overlapping edges between HBN and PNC shown as the bold value in parentheses; the second percentage shows the minimum percentage of edges required for results to be significant. The shaded regions in each circular bar show the minimum percentage of edges required for significance with a p-value below 0.05. * above each of the labels show significance for each network and the whole brain. *indicates p ≤ 0.05, **indicates p ≤ 0.01, and ***indicates p ≤ 0.001

**Fig. S10:**
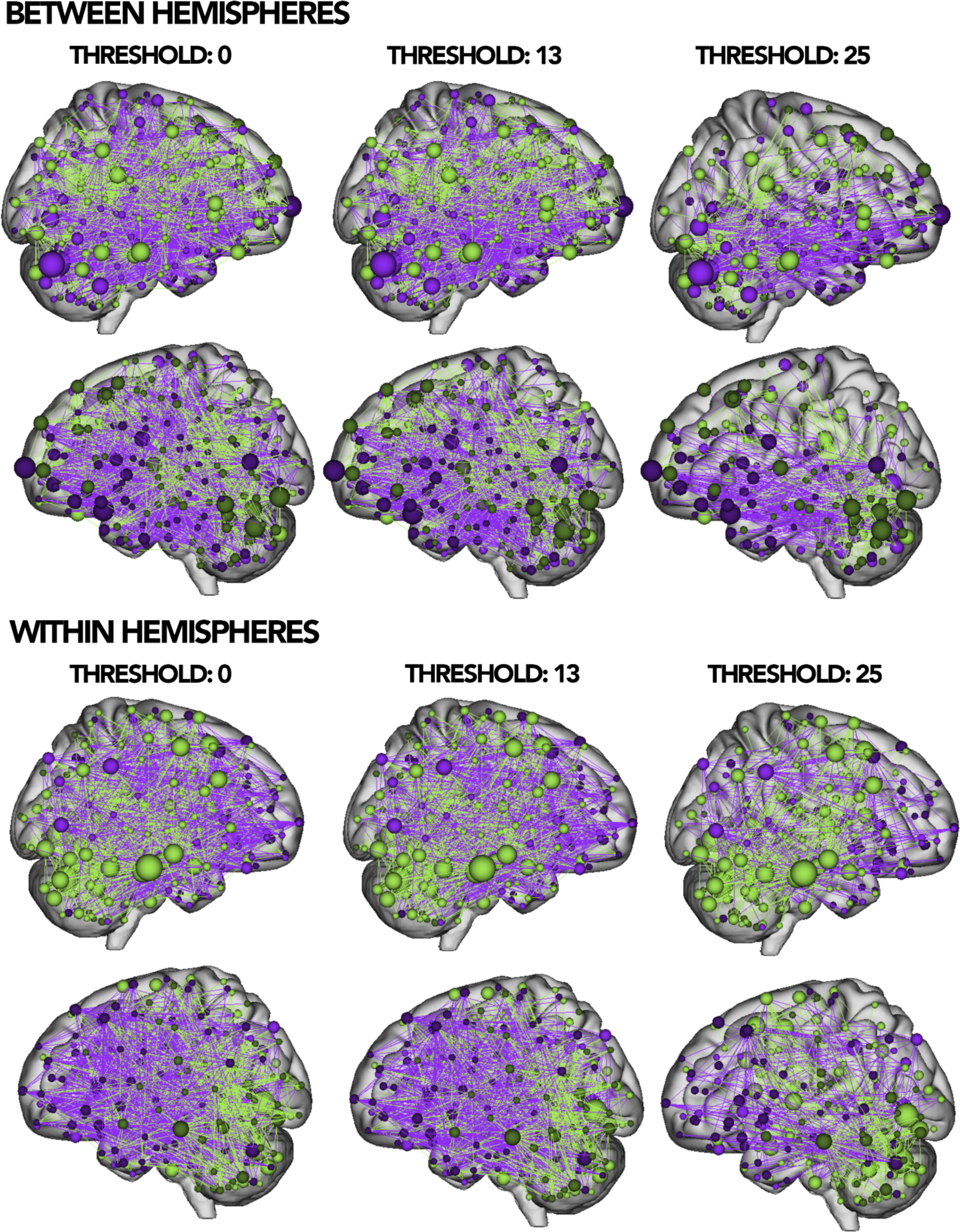
Brain plots for differences between left-handed individuals and right-handed individuals between and within-hemispheres for combined analyses with HBN and PNC thresholded at degree = 0, 13, and 25.

**Fig. S11:**
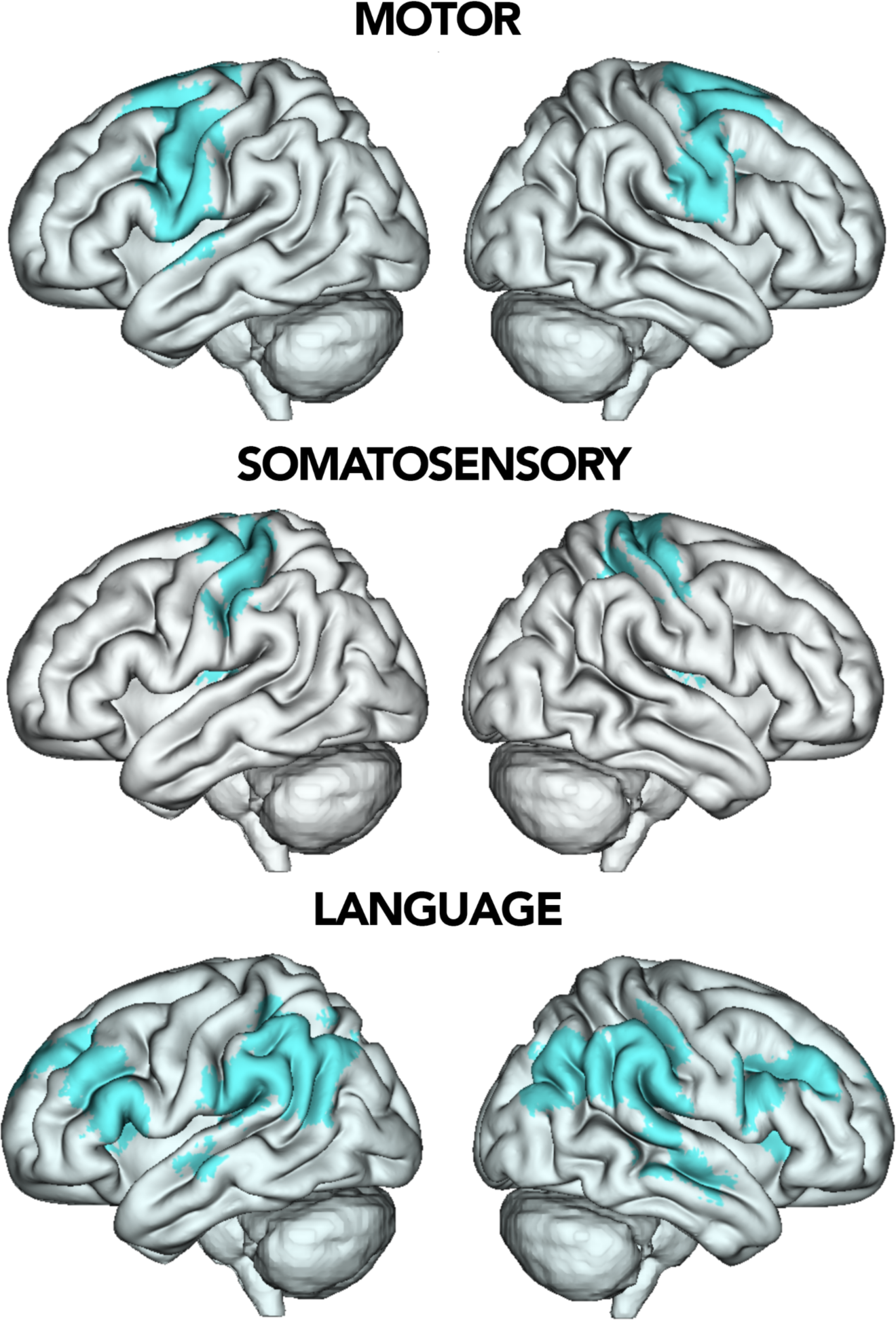
3D brain plots showing where nodes for each network of interest are located.

**Fig. S12:**
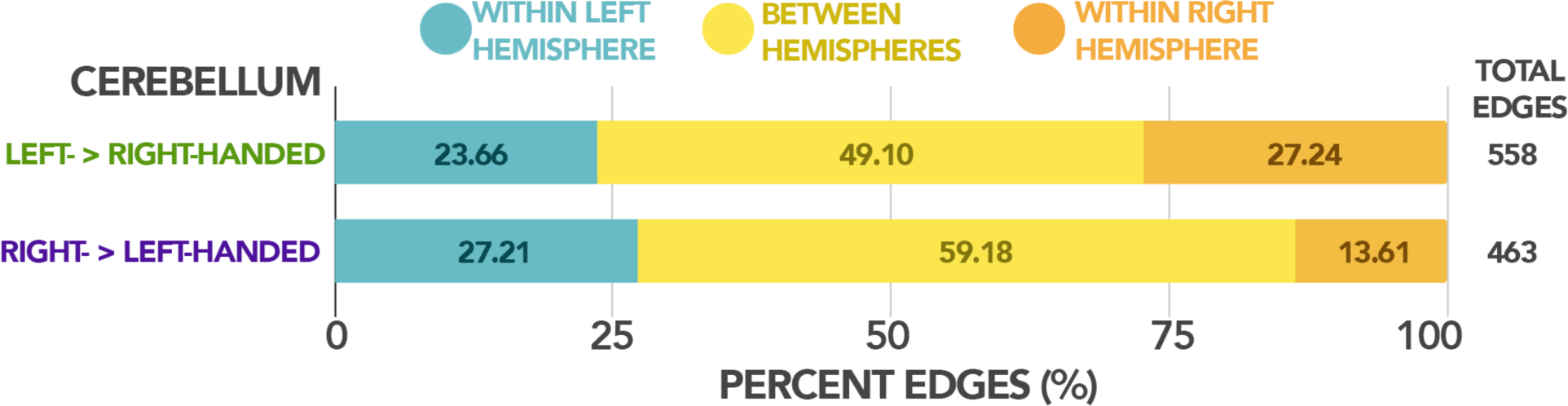
Percent of edges in our cerebellum analysis split by left- and right-handed individuals connecting within left/right hemispheres or between-hemispheres.

### Supplementary Tables

**Table S1:** Replicating all HBN analyses in networks of interest: motor, somatosensory, language, and HBN+PNC analyses in whole brain and cerebellum while controlling for sex. Table shows comparison between significant edges when controlling for sex and for handedness alone, the overlapping edges between these two analyses and the correlation between the two analyses (reported as r values).

|  | Handedness group | Control for Sex | Handedness only | Overlapping edges | r-val |
| --- | --- | --- | --- | --- | --- |
| **HBN: Motor** | **Left > Right** | 194 | 195 | 181 | 0.9917 |
|  | **Right > Left** | 217 | 227 | 205 | 0.9879 |
| **HBN: Somatosensory** | **Left > Right** | 93 | 88 | 85 | 0.9912 |
|  | **Right > Left** | 105 | 112 | 100 | 0.9903 |
| **HBN: Language** | **Left > Right** | 338 | 325 | 315 | 0.9937 |
|  | **Right > Left** | 332 | 337 | 324 | 0.9943 |
| **HBN+PNC: Whole Brain** | **Left > Right** | 1566 | 1600 | 1501 | 0.9888 |
|  | **Right > Left** | 1452 | 1479 | 1396 | 0.9865 |
| **HBN+PNC: Cerebellum** | **Left > Right** | 529 | 558 | 509 | 0.9938 |
|  | **Right > Left** | 439 | 463 | 431 | 0.9940 |

**Table S2:**
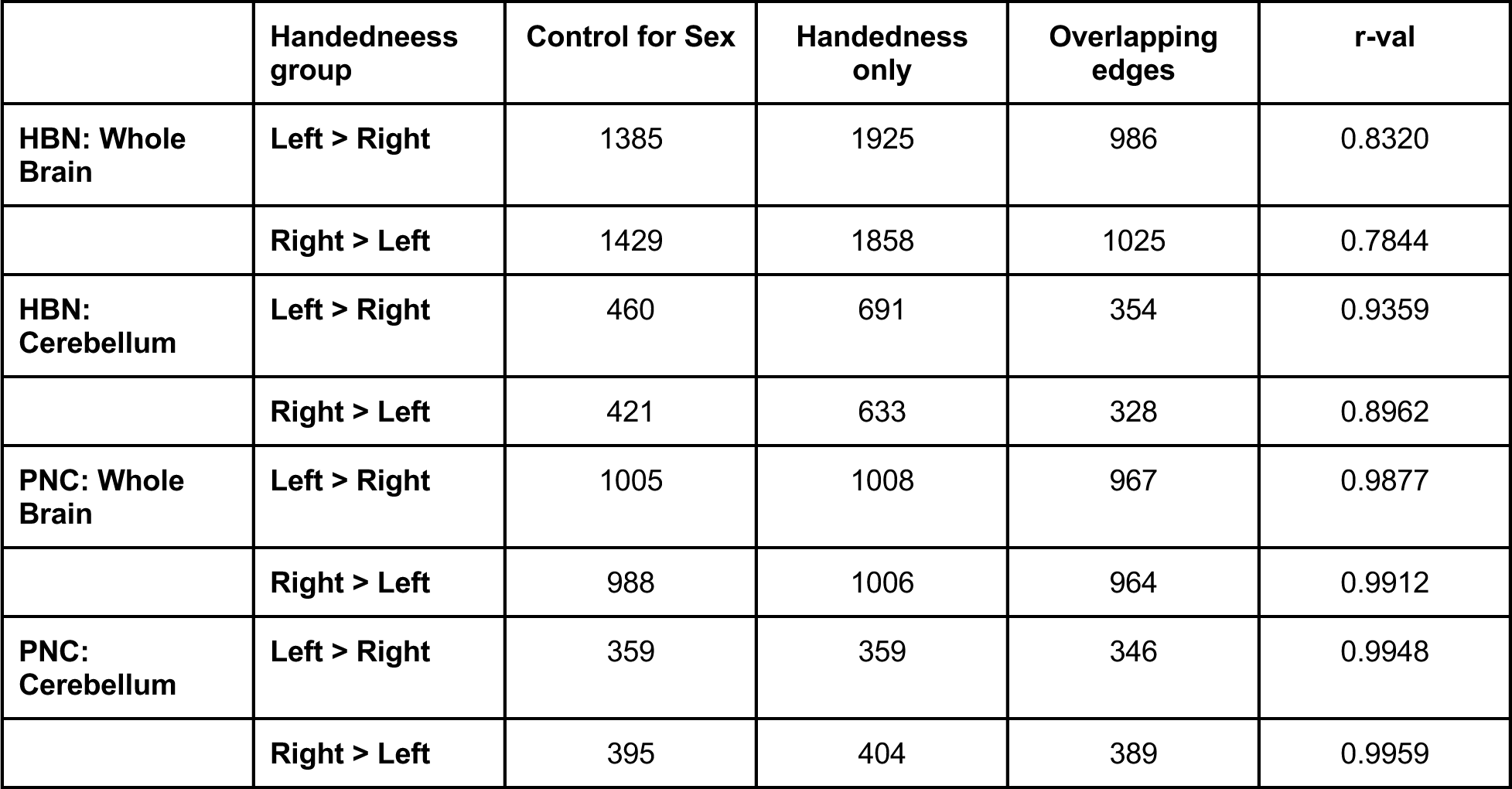
Conducting whole brain and cerebellum analyses for HBN and PNC. Comparing significant edges while controlling for sex and for handedness alone, the overlapping edges between these two analyses and the correlation between the two analyses (reported as r values).

**Table S3:** Replicating all HBN analyses in networks of interest: motor, somatosensory, language, and HBN+PNC analyses in whole brain and cerebellum while controlling for age. Table shows comparison between significant edges when controlling for age and for handedness alone, the overlapping edges between these two analyses and the correlation between the two analyses (reported as r values).

|  | Handedness group | Control for Age | Handedness only | Overlapping edges | r-val |
| --- | --- | --- | --- | --- | --- |
| HBN: Motor | Left > Right | 149 | 195 | 147 | 0.9620 |
|  | Right > Left | 187 | 227 | 179 | 0.9800 |
| HBN: Somatosensory | Left > Right | 62 | 88 | 62 | 0.9660 |
|  | Right > Left | 91 | 112 | 86 | 0.9748 |
| HBN: Language | Left > Right | 252 | 325 | 245 | 0.9809 |
|  | Right > Left | 269 | 337 | 262 | 0.9710 |
| HBN+PNC: Whole Brain | Left > Right | 8382 | 1600 | 749 | 0.9010 |
|  | Right > Left | 7872 | 1479 | 682 | 0.8982 |
| HBN+PNC: Cerebellum | Left > Right | 4014 | 558 | 235 | 0.6406 |
|  | Right > Left | 3669 | 463 | 206 | 0.5272 |

**Table S4:** Conducting whole brain and cerebellum analyses for HBN and PNC. Comparing significant edges while controlling for age and for handedness alone, the overlapping edges between these two analyses and the correlation between the two analyses (reported as r values).

|  | Handedness group | Control for Age | Handedness only | Overlapping edges | r-val |
| --- | --- | --- | --- | --- | --- |
| HBN: Whole Brain | Left > Right | 1526 | 1925 | 1153 | 0.7941 |
|  | Right > Left | 1473 | 1858 | 1199 | 0.7104 |
| HBN: Cerebellum | Left > Right | 537 | 691 | 376 | 0.9179 |
|  | Right > Left | 485 | 633 | 327 | 0.8508 |
| PNC: Whole Brain | Left > Right | 981 | 1008 | 938 | 0.9800 |
|  | Right > Left | 953 | 1006 | 932 | 0.9833 |
| PNC: Cerebellum | Left > Right | 349 | 359 | 336 | 0.9914 |
|  | Right > Left | 368 | 404 | 358 | 0.9906 |

**Table S5:**
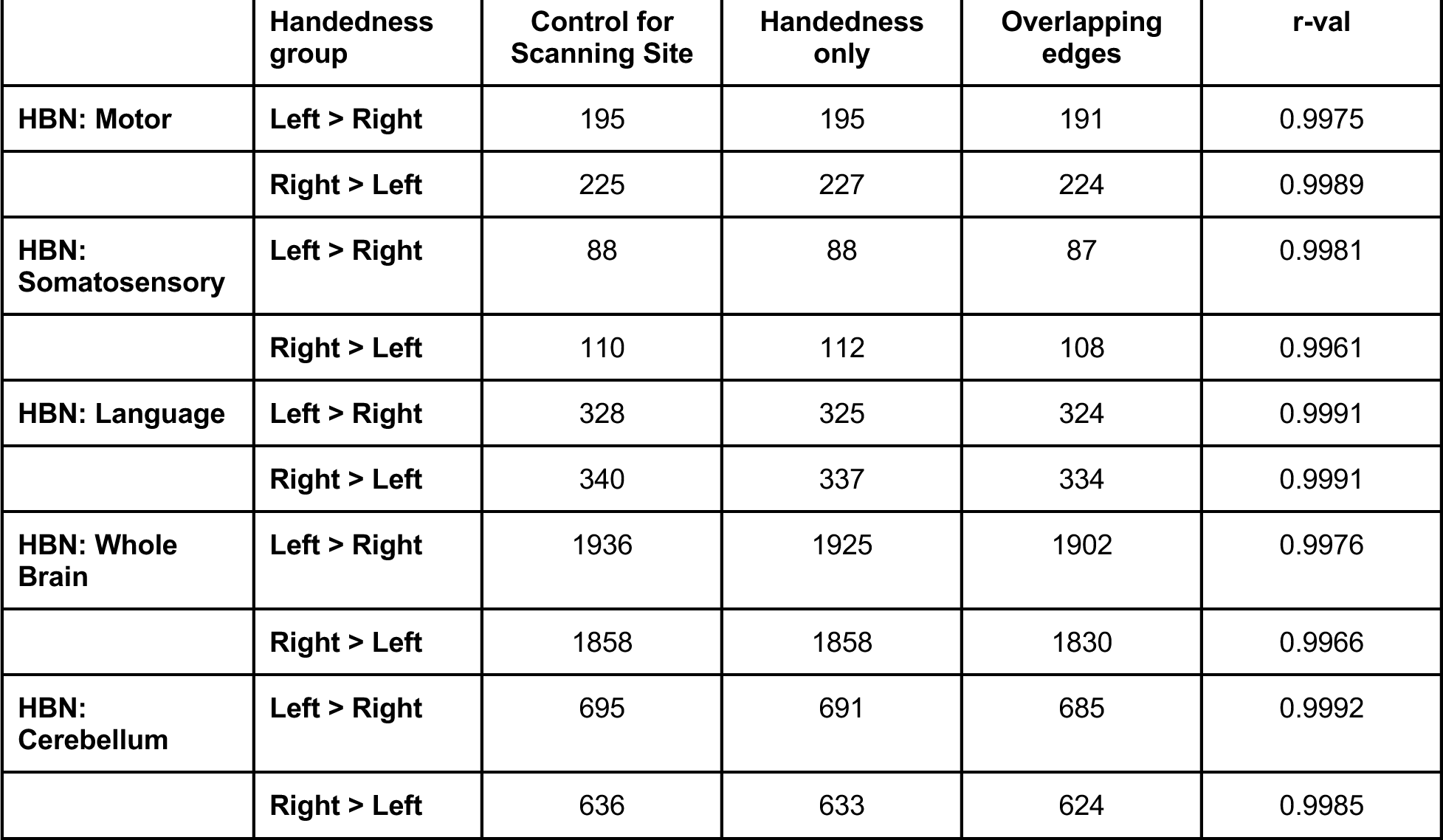
Replicating all HBN analyses in networks of interest: motor, somatosensory, language, as well as analyses in whole brain and cerebellum while controlling for scanning site. Table shows comparison between significant edges when controlling for scanning site, the overlapping edges between these two analyses and the correlation between the two analyses (reported as r values).

**Table S6:** Replicating all HBN analyses in networks of interest: motor, somatosensory, language, as well as analyses in whole brain and cerebellum while controlling for clinical diagnoses. Table shows comparison between significant edges when controlling for clinical diagnoses, the overlapping edges between these two analyses and the correlation between the two analyses (reported as r values).

|  | Handedness group | Control for Clinical Diagnoses | Handedness only | Overlapping edges | r-val |
| --- | --- | --- | --- | --- | --- |
| HBN: Motor | Left > Right | 195 | 195 | 191 | 0.9975 |
|  | Right > Left | 226 | 227 | 224 | 0.9989 |
| HBN: Somatosensory | Left > Right | 88 | 88 | 87 | 0.9988 |
|  | Right > Left | 110 | 112 | 108 | 0.9961 |
| HBN: Language | Left > Right | 328 | 325 | 324 | 0.9991 |
|  | Right > Left | 340 | 337 | 336 | 0.9980 |
| HBN: Whole Brain | Left > Right | 1936 | 1925 | 1902 | 0.9976 |
|  | Right > Left | 1858 | 1858 | 1830 | 0.9966 |
| HBN: | Left > Right | 695 | 691 | 685 | 0.9992 |
| Cerebellum |  |  |  |  |  |
|  | Right > Left | 636 | 633 | 624 | 0.9985 |

